# Substrate recognition and cleavage by mucin degrading *O*-glycopeptidases from the gut microbe *Bacteroides caccae*

**DOI:** 10.1101/2025.10.14.682407

**Authors:** Benjamin Pluvinage, Kathleen Bourdon, Olivia Canil, Ashley Deventer, Bernadette Alvarez, Liam Mihalynuk, Nicole Thompson, Warren Wakarchuk, Alisdair B. Boraston

**Affiliations:** Department of Biochemistry and Microbiology, University of Victoria, PO Box 1700 STN CSC, Victoria, British Columbia, V8W 2Y2, Canada; Department of Biological Sciences, University of Alberta, Edmonton, AB T6G 2E9, Canada; Biomedical Sciences Research Complex, School of Biology, University of St Andrews, St Andrews, UK

## Abstract

*O*-glycopeptidases are enzymes that hydrolyze the peptide bonds in glycoproteins by a mechanism that involves specific recognition of *O*-linked glycans on the substrate. *Bacteroides caccae*, an accomplished mucin degrader, is a member of the human gut microbiota with sixteen genes encoding putative *O*-glycopeptidases in the peptidase_M60 family. At present, the diversity of substrate selective in *O*- glycopeptidases is not well-understood nor is the rationale behind their expansion in bacteria such as *B. caccae*. Here we reveal the activity and diversity of the peptidase_M60 *O*-glycopeptidases encoded in the *B. caccae* genome. At least thirteen of the sixteen peptidase_M60 genes are active mucinases. Targeted functional studies by a high-throughput FRET screen combined with detailed kinetic analyses reveal that five examples in an uncharacterized clade of peptidase_M60 proteins have different substrate selectivities despite their relatively high degree of relatedness. Structural analyses of these enzymes, including bound complexes, reveal new insight into the molecular underpinnings of *O*-glycopeptidase diversity. This highlights the larger context of how varied the selectivity of peptidase_M60 *O*-glycopeptidases can be for the glycan moiety and/or the peptide portion of the substrates, and why mucin degraders like *B. caccae* diversify *O*-glycopeptidase substrate repertoires to potentially maximize breakdown of this extraordinarily complex polymer.

## INTRODUCTION

As one of the first barriers a pathogen encounters when entering the body, mucus plays a key role in host defense ^1^. Mucosal surfaces in the human body, such as in the nose, gut, and, mouth, utilize a thick viscous substance to lubricate and protect its vulnerable entryways ^2^. The main components of mucus are glycoprotein subunits called mucins that are produced by epithelial cells and comprised of two main parts: a peptide backbone with complex, branched carbohydrate chains ^3^. Over 80% of mucus mass comes from the carbohydrate portion, which are mostly *O*-linked glycans. Mucins are most densely glycosylated at stretches of the peptide backbone containing multiple proline, threonine, or serine residues (PTS domains), that are present in varying numbers of tandem repeats ^4,5^. The large amounts of *O*-linked glycans extending from mucins associate with water and protect the peptide backbone from general protease cleavage, two features that make mucus an ideal protector for bodily surfaces that come in contact with microbes ^4,6^.

Colonic epithelial cells have a dense layer of mucus that prevents microbiota residents from infiltrating and this layer is considered sterile in absence of infection ^3^. This mucosal layer becomes thinner and more loosely packed the further into the lumen it extends. Both commensal and pathogenic microbes occupy the outer mucosal layer by utilizing a suite of enzymes to break down mucins ^7,8^. The outer mucus layer is turned over rapidly: the processes of mucin degradation by the microbiota and mucin production from the secretory epithelial cells exist in an equilibrium to maintain homeostasis. Mucin-degrading enzymes can be upregulated in times when the amount of dietary polysaccharides are not sufficient to provide the microbe with enough energy ^9^. These specialized enzymes, which include glycosidases, give some microbes a competitive edge over non-mucin-degrading bacteria, by providing unique mechanisms for adapting to mucosal environments ^7,10,11^. The microbiota comprises a diverse range of commensal and mutualistic microbes, and the capabilities of these microbes to degrade mucin have not yet been fully revealed.

Recently, members of the enzyme superfamily peptidase_M60 (Pfam13402) were found to be widely distributed in the genomes of many host-adapted microbes ^12^. Examples of this domain were revealed to be a new type of metallopeptidase with specificity for *O*- glycosylated substrates, such as mucins ^13^. These zinc-dependent enzymes are endo-acting peptidases on the N-terminal side of an *O*-glycosylated residue ^13–15^. Deemed *O*- glycopeptidases, these domains require an *O*-linked GalNAc for activity, with the ability to tolerate more complex glycans in enzyme subsites ^13^. Glycan-binding sites, or G-sites, that can accommodate different glycan groups were proposed to be the main strategy to determine recognition of glycans, and thus enzyme specificity. Conversely, the peptide sequence of the substrate has an as-yet undefined impact on *O*-glycopeptidase substrate specificity. The unique activity of the peptidase_M60 *O*-glycopeptidases on *O*- glycosylated substrates, such as mucins, make them key candidates for the interaction of microbes with mucins and important tools used in mucin foraging strategies.

The genus Bacteroides accounts for a large portion of the gut microbiota with some species known to have the capacity to degrade mucin ^1,2,16^. *B. caccae* is notable in this respect ^9^. Its genome encodes numerous degradative glycosidases with known or predicted activities associated with host-glycan degradation. It also encodes sixteen putative peptidase_60 *O*-glycopeptidases ^9^, fifteen of which are associated with polysaccharide utilization loci (PUL)^17^. All but one of the genes encoding these proteins showed increased *in vivo* expression under dietary fibre deprivation in a mouse model of colonic colonization ^9^. This suggested a role in mucin foraging; however, the activities of the putative *B. caccae O*-glycopeptidases and why the genes encoding these are so expanded in this microbe remain unknown. Here we begin to unravel this by demonstrating the mucinase activity of twelve of the putative *B. caccae O*- glycopeptidases. Furthermore, we target five examples representing sub-clades of an uncharacterized peptidase_M60 clade and reveal the molecular details of their specificities using a novel FRET-based screen, kinetic analyses, and structural studies. The results reveal many of the molecular underpinnings of the processes *B. caccae* uses to process the backbone mucins that separate host tissues from this colonizing bacterium.

## RESULTS

### Bacteroides caccae O-glycopeptidases

We initially assessed the diversity of the *B. caccae* peptidase_M60 metallopeptidase domains by comparing their amino acid sequences **(Figure 1)**. These sequences displayed pairwise identities as low at 21% and up to 74%. Two clear clusters of sequences emerged with a third split cluster and then BcM60G as a relative outlier. To place these sequences in the greater context of peptidase_M60 uperfamily we extracted the sequences of bacterial origin from Pfam entry 13402 (Pfam version 32). A multiple sequence alignment of the truncated metallopeptidase domain sequences from >500 unique entries, which we ensured included the sixteen *B. caccae* proteins, revealed a maximum sequence identity of ∼93% and a minimum of ∼15%. The protein phylogeny constructed from this alignment placed in the context of known *O*-glycopeptidases reveals the enormous sequence space of the family and how little of it has been sampled by enzyme characterization **(Supplementary Figure 1)**. The *B. caccae* proteins fell into four separate clades in the overall analysis, consistent with their clustering in the pairwise comparison of only the *B. caccae* proteins. BcM60A and BcM60E, however, were satellites to Clade 3 showing their pairwise separation from BcM60D and BcM60H. BcM60G was the lone *B. caccae* entry in Clade 2.

**Figure 1.**
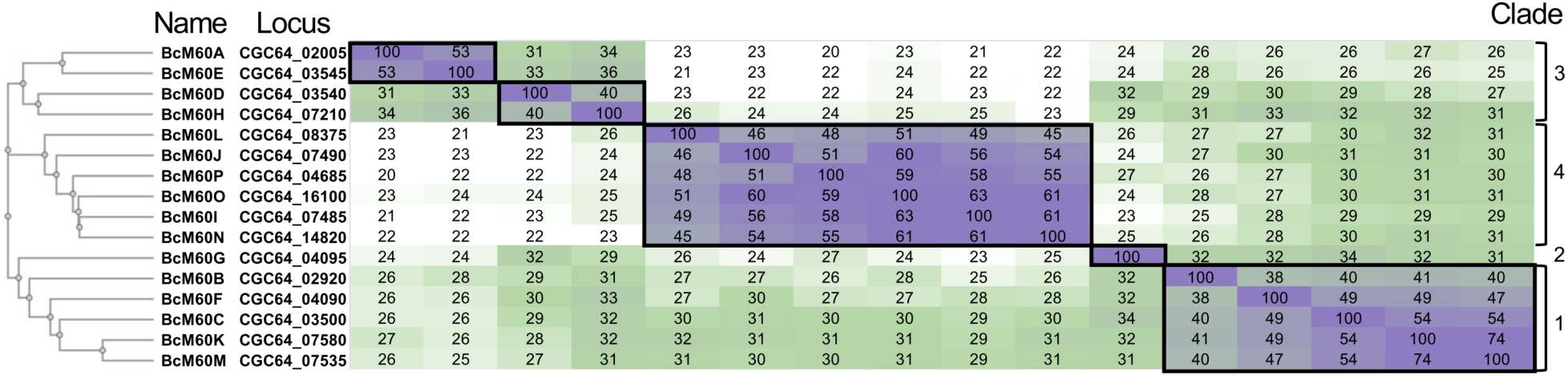
Comparison of the Bacteroides caccae peptidase_M60 proteins. Percent amino acid sequence identity matrix for the sixteen peptidase_M60 proteins from *B. caccae*.

We recombinantly produced and purified the metallopeptidase domains of twelve *B. caccae O*-glycopeptidases that spanned the sequence diversity of the *B. caccae* enzymes. For brevity, these truncated constructs are referred to as BcM60A to BcM60L. These proteins all showed activity in a microplate-based mucinase activity, and inhibition by EDTA, consistent with metallopeptidase mucinolytic activity **(Figure 2)**. Though the results with BcM60B were statistically significant it did appear to be the least active of the twelve enzymes.

**Figure 2.**
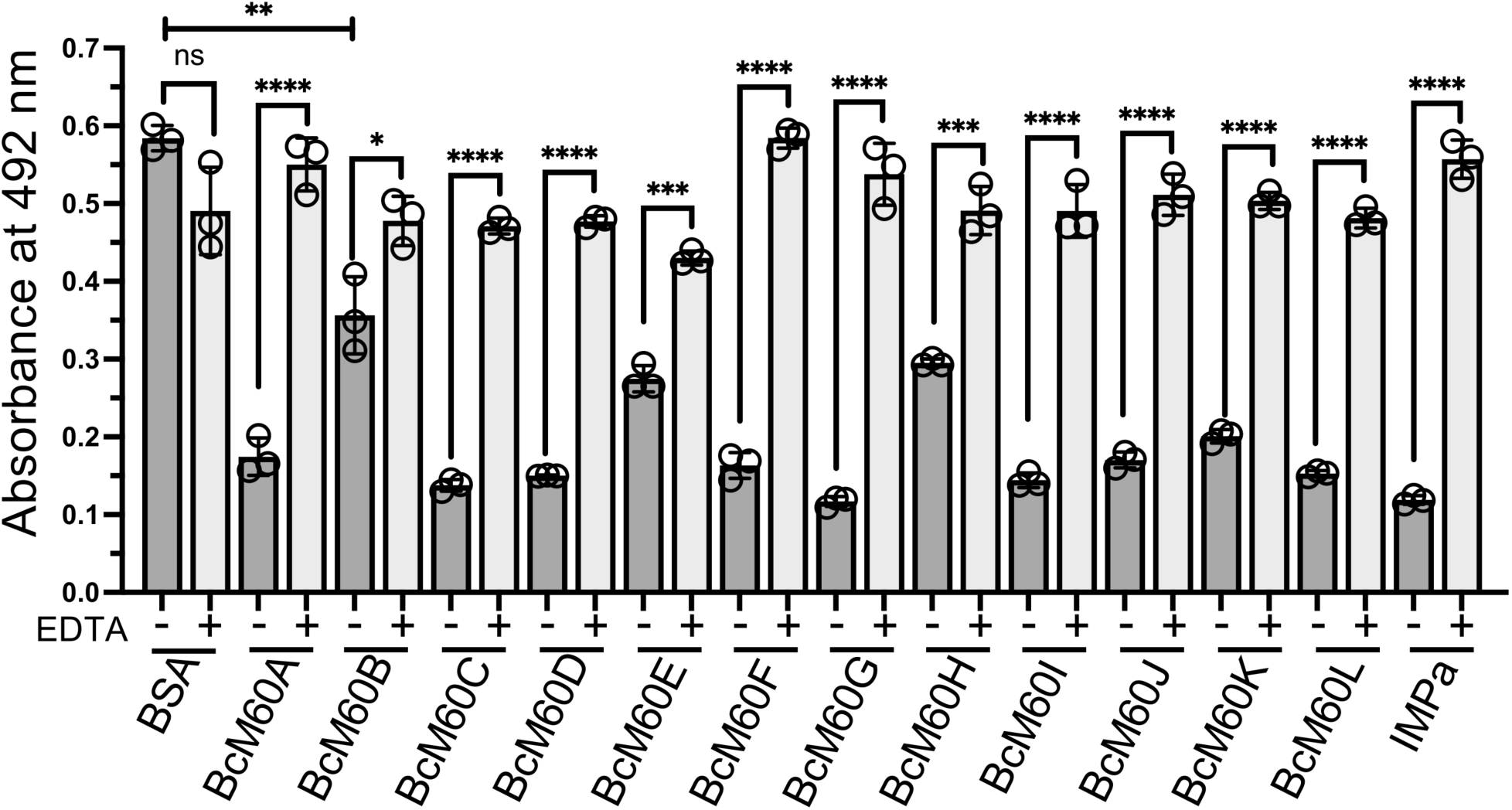
Mucinase assay showing the activity of twelve *B. caccae* peptidase_M60 proteins on bovine maxillary mucin (BSM). A reduction in signal indicates loss of immobilized biotinylated BSM and therefore mucinase assay. Pairwise comparisons of the no-EDTA samples to the BSA control all gave P-values <0.0001, except for BM60B, which is indicated in the figure. Pairwise comparisons of the EDTA samples to the BSA EDTA control sample all gave P-values > 0.05. IMPa was included as a positive control. P-value symbols for unpaired two-tailed T-tests are as follows: P<0.05 (*), P<0.01 (**), P<0.001 (***), P<0.0001 (****).

To expand our knowledge of the M60-like *O*-glycopeptidase sequence space we targeted the large clade 1, which comprises up to three distinct sub-clades and lacks any characterized members. We used the recombinant constructs truncated to the metallopeptidase domains of BcM60B, BcM60C, BcM60F, BcM60K, and BcM60M to represent all *B. caccae* members of this clade.

### Substrate screening

To rapidly provide a semi-quantitative assessment of *O*- glycopeptidase activity we built a high-throughput screen. The substrates for our fluorescent protein-based FRET screen for *O*-glycopeptidases were closely based on those we used previously and comprised mNeonGreen at the N-terminus and mScarlet-I at the C-terminus ^18^. An eleven amino acid linker separated the fluorescent protein domains. Twenty-four PTS repeats representing fourteen common human mucins and three non-mucin proteins were chosen to comprise the linker sequences (**Supplementary Table 1**). The linker sequences were then glycosylated to bear Tn (α- linked O-GalNAc), C1 (Galβ1-3GalNAcα-), 3SC1 (Neu5Acα2-3Galβ1-3GalNAcα-), 6SC1 (Galβ1-3[Neu5Acα2-6]GalNAcα-) and or C2 (Galβ1-3[GlcNAcβ1-6]GalNAcα-). Each glycan lineage of substrate was prepared from one large preparation of the Tn modified protein. The C1, C2 and 6SC1 versions of the MUC2_R1, MUC5AC_R2, IgA1 hinge, and MUC16 modified substrates were analyzed by mass spectrometry (**Supplementary Figure 2**). This revealed that only a small proportion of the substrates remained unglycosylated and conversion of Tn to C1 was essentially complete. The C2 substrates were fully converted in all the substrates. Only the MUC5AC_R2 linker was fully converted to 6SC1 with the other three having residual C1 modified substrate. The MUC5AC_R2 and IgA1 hinge linkers each only had a single glycan, presumably at one site, though the IgA1 linker has several predicted acceptor sites in addition to the main one. The MUC2_R1 and MUC16 linkers had 1-4 and 1-2 glycans, respectively, indicating heterogeneity as well as microheterogeneity in the 6SC1 samples.

Using BT4244 as our benchmark enzyme, which was the founding peptidase_M60 *O*- glycopeptidase ^12^, we optimized the excitation wavelength and emission wavelengths for mNeonGreen and mScarlet-I, as we did previously ^18^. As with the prior substrates, an excitation wavelength of 430 nm was found to provide optimal emission from mNeonGreen at 518 nm without significant direct excitation of mScarlet-I. FRET emission from mScarlet-I in intact substrate was found to be optimum at 590 nm.

With 47 different substrates containing varied linkers and glycans including Tn, C1, C2, 3SC1, and 6SC1 we tested our screen with BT4244. Notably, BT4244 displayed high activity with the FRET signal, determined as a “ratio-of-ratios” (RoR)^18^, often hitting a maximum at between 1-4 hours of reaction time (**Figure 3**). BT4244 had broad activity on 17 of the 24 linkers, though it had qualitatively lower activity on the IgA1 hinge repeat as revealed by the extended time necessary to produce a signal for substrate cleavage. Consistent with prior studies, BT4244 had activity on substrates with the Tn, C1, and 3SC1 glycans. BT4244 lacked activity on substrates with C2 glycans, indicating intolerance of branched glycans. It had some activity on MUC2_R1 and MUC16 with the 6SC1 glycan, but these are known to have some degree of residual C1 leading us to suspect the signal was from cleavage of this contaminating substrate. Supporting this is the observation that BT4244 had good activity on the MUC5AC_R2 linker with Tn, C1, and 3SC1 glycans but no detectable activity on this linker with 6SC1, which was essentially completely modified and thereby lacked contaminating C1. This screen was thus successful in providing a rapid readout of BT4244 activity that was consistent with prior knowledge of this enzyme ^12–14,18^.

**Figure 3.**
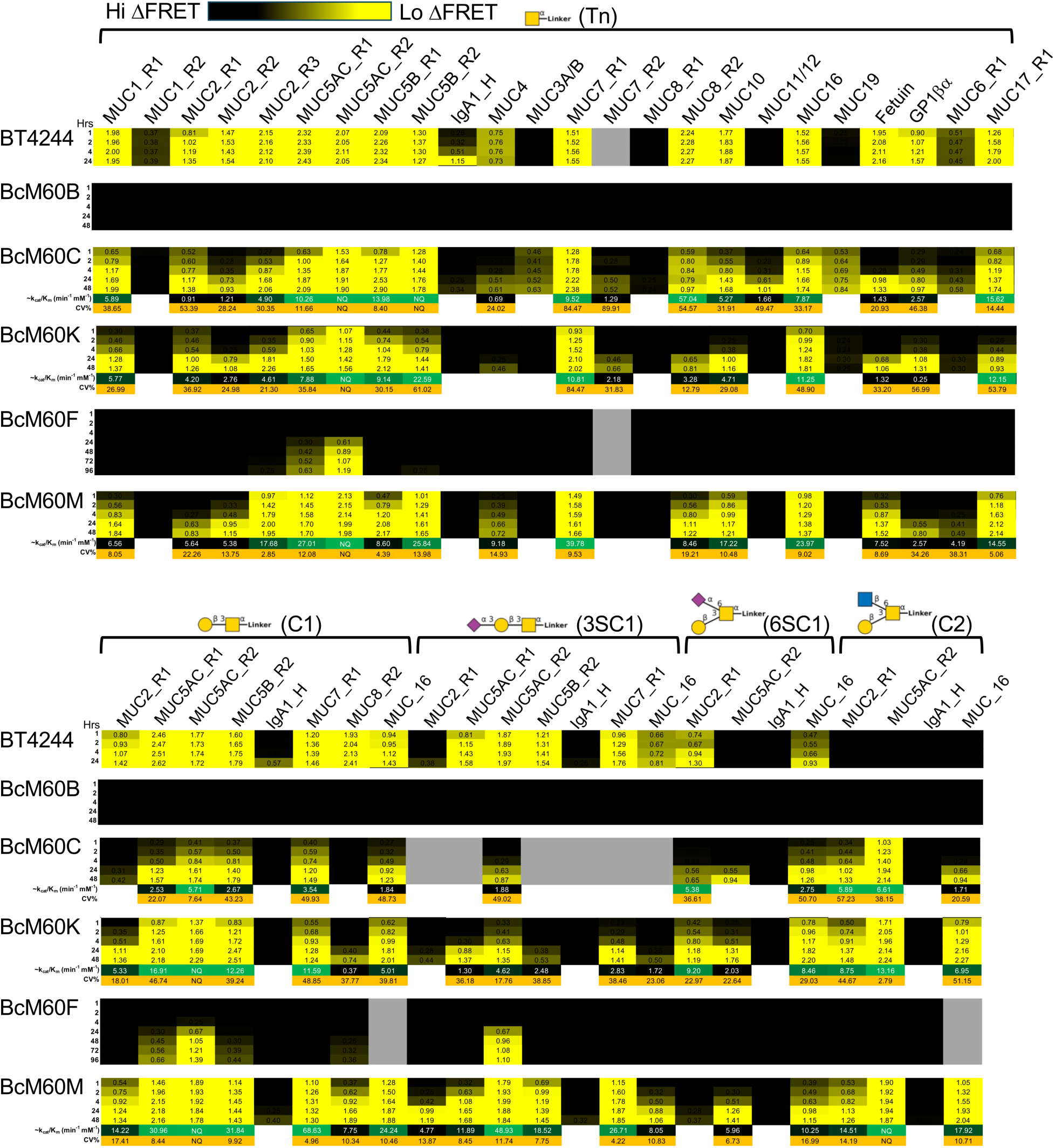
*O*-glycopeptidase activity revealed by a custom screen of mucin-like FRET substrates. Heat maps show FRET values calculated by the RoR approach. The coloring is ramped from a low threshold of 0.2 to a high of 1.0. Where possible, k_cat_/K_M_ were determined by fitting FRET signal as fluorescence difference values with a simple E + S → E + P model. These values, along with the coefficient of variance to estimate confidence, are shown below the heatmap.

We then applied this screen to BcM60B, BcM60C, BcM60F, BcM60K and BcM60M (**Figure 3**). Despite observing low activity in the mucinase assay for BcM60B, we failed to detect activity in the FRET screen, possibly indicating a unique recognition determinant not represented in the screen. BcM60C, BcM60K, and BcM60M displayed nearly identical fingerprints of substrate selectivity, but unlike BT4244, they had activity on C2 substrates and the MUC5AC_R2 6SC1 substrate, indicating accommodation of these branched *O*- glycans. The three enzymes appeared qualitatively less active than BT4244, with BcM60C apparently being the least active as it had to be assayed at ∼2 times the enzyme concentration to generate similar signal. BcM60M was assayed at the same concentration as BcM60C and, qualitatively, showed greater activity. None of the enzymes were active on non-glycosylated versions of the substrates (**Supplementary Figure 3**)

For those enzyme-substrate combinations showing significant activity by the RoR we constructed progress curves using fluorescence difference values and analyzed them using a simplified bi-molecular model of E + S ←→ E + P to estimate k_cat_/K_M_ values, an approach sometimes referred to as “hit and run” (see **Supplementary Figure 4**). Fits were rejected if they were visibly nonsensical and/or the residuals failed a statistical “runs” test. In some cases, such as with BT4244, some of the reactions progressed too quickly to perform this analysis. This analysis added value over the semi-quantitative visual interpretation of the RoRs by enhancing assessment of relative enzyme efficiency within a screen. Even with this analysis there was nothing significant to distinguish between the substrate selectivity of BcM60K, BcM60M, and BcM60C, though the latter enzyme appears to be slightly less efficient. However, this clearly revealed a 5-10 fold reduced activity on the 3SC1 substrates for both enzymes. BcM60M appeared to have only a 2-3 fold reduction in activity on 3SC1 modified substrates compared to the C1 substrates, suggesting it may be more tolerant of this modification, but had relatively poor activity on the MUC5AC_R2 6SC1 substrate.

We further explored the influence of a terminal α-2,3-linked sialic acid using BcM60K. We did this by comparing activity on 3SC1 substrates and 3SC1 substrates co-treated with NanH, a highly active α-2,3-sialidase ^19^. The RoR analysis revealed a clear increase in activity in the presence of NanH. The hit and run analysis also showed higher rates of conversion in the presence of NanH with 3-5 fold increases in the estimates of k_cat_/K_M_ (**Figure 4**).

**Figure 4.**
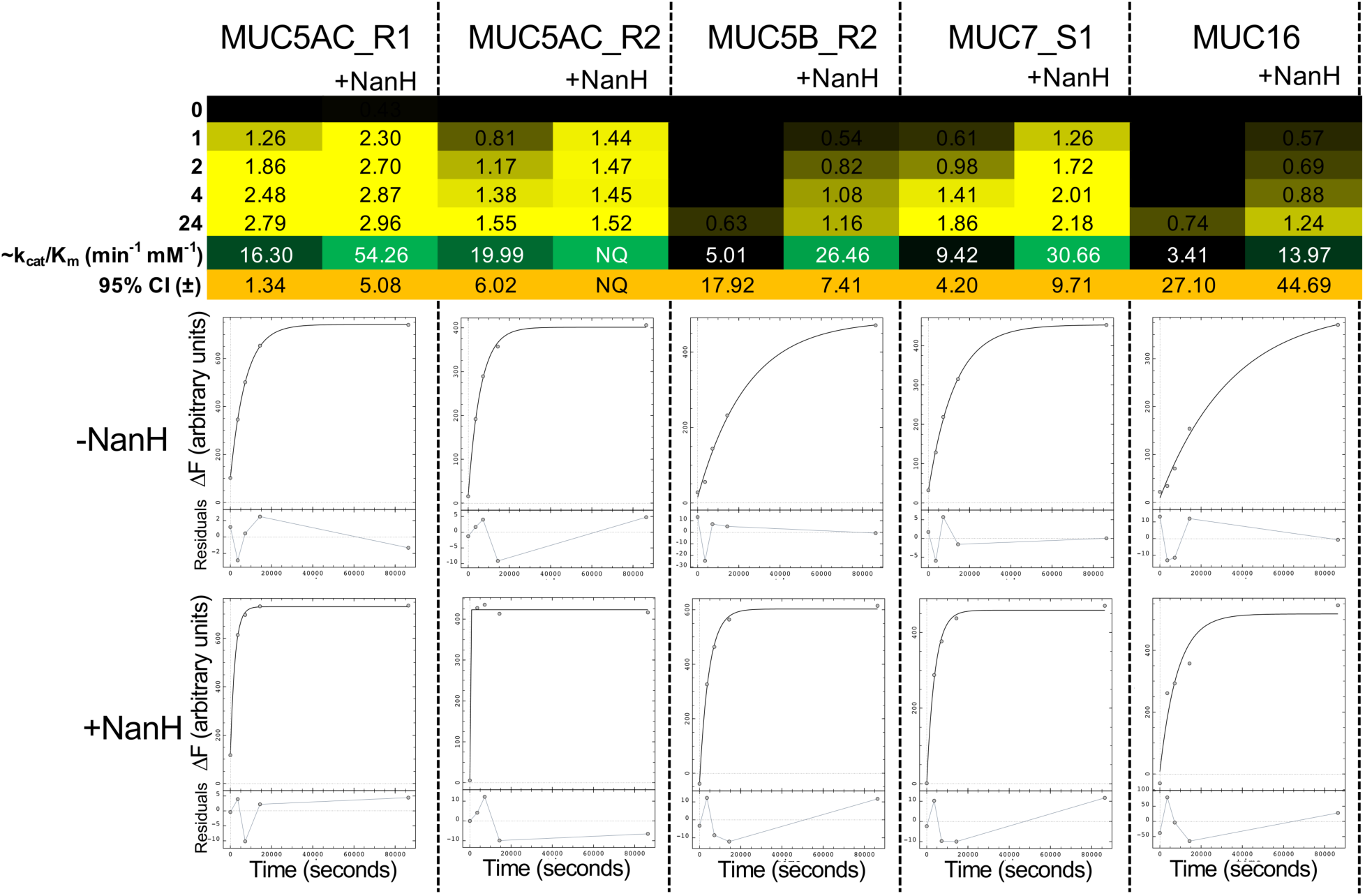
NanH potentiates BcM60K activity on substrates with 3SC1 glycans. The FRET screen based on RoR analysis is shown for experiments done in the presence or absence of NanH. Values below the heat map show the results of fits of the difference fluorescence analysis with the actual curves and fits shown below for reference.

### Quantitative analysis of glycan selectivity

The screen revealed clear selectivity of the enzymes for certain glycans and peptide sequences. We explored this quantitatively using BcM60C, BcM0K, BcM60F, and BcM60M with the FRET assay adapted to kinetic analysis, which employed a global analysis of multiple progress curves to enable determination of detailed kinetic parameters (see **Supplementary Figure 5**). We focused on substrates for which we had detailed mass spectrometric analysis of the glycosylation state (MUC2_R1, MUC5AC_R2, and/or IgA1_h). In this analysis, though a small amount of residual unglycosylated material was present, we assumed complete cleavage. Indeed, when modelling the kinetics with 10-20% uncleavable substrate the results obtained were within the 95% confidence intervals when assuming complete cleavage.

We again used BT4244 as our benchmark for the assay and enzyme activity. Single digit k_cat_ values per minute revealed generally slow turnover while single digit μM K_M_ values indicated a high apparent affinity. The enzyme efficiencies (k_cat_/K_M_) varied approximately 3-fold with the MUC2_R1 and IgA1_h linkers being the less preferred substrate and the MUC5AC_R2 linker the most favored. The k_cat_/K_M_ values for the MUC2_R1 and IgA1_h linkers was similar to the value of 594 min^-1^ mM^-1^ obtained for an engineered *O*- glycosylation sequon in an FRET substrate with nearly identical architecture, albeit determined by a slightly different approach that only yielded k_cat_/K_M_ ^18^.

The k_cat_ values for BcM60K also indicated single digit catalytic events per minute, like BT4244 **(Table 1)**. Likewise, K_M_ values also revealed high apparent affinity for substrate. However, these values combined to give catalytic efficiencies that were about 2-4 fold lower than those for BT4244. The pattern of substrate selectivity was, from preferred to least preferred, C1 > Tn ≈ 6SC1 > 3SC1 ≈ C2 with an ∼4-fold difference between the efficiencies of the most and least preferred substrates. The fold difference between the C1 and 3SC1 substrates by this detailed analysis was consistent with that observed by the analysis of the FRET screen with and without NanH, indicating the clear detrimental influence of the terminal α-2,3-linked sialic. BcM60C preferred the C1 substrate over the Tn with catalytic efficiencies slightly below half that of BcM60K **(Table 1)**, which is consistent with reduced activity in the FRET screen.

**Table 1:** Kinetic parameters.

| Enzyme | Linker | O-glycan | $k_{cat}/K_M$<br>($\text{min}^{-1} \text{mM}^{-1}$) | $k_{cat}$ ($\text{min}^{-1}$) | $K_M$ ($\mu\text{M}$) |
| --- | --- | --- | --- | --- | --- |
| BT4244 | MUC5AC_R2 | Tn | 2190.2 (± 166.6) | 6.3 (± 0.6) | 2.9 (± 0.1) |
| | | C1 | 2182.5 (± 193.4) | 6.3 (± 0.7) | 2.9 (± 0.1) |
| | MUC2_R1 | Tn | 751.6 (± 36.1) | 4.2 (± 0.2) | 5.6 (± 0.1) |
| | IgA1_hinge | Tn | 871.2 (± 198.5) | 0.9 (± 0.1) | 1.0 (± 0.1) |
| BcM60K | MUC5AC_R2 | Tn | 443.4 (± 22.8) | 3.4 (± 0.4) | 7.8 (± 1.0) |
| | | C1 | 556.6 (± 24.8) | 3.4 (± 0.4) | 6.2 (± 0.7) |
| | | 3SC1 | 133.5 (± 9.5) | 1.0 (± 0.1) | 7.6 (± 1.0) |
| | | 6SC1 | 92.6 (± 30.9) | 0.2 (± 0.1) | 2.5 (± 0.7) |
| | | C2 | 401.5 (± 21.0) | 1.8 (± 0.2) | 4.5 (± 0.4) |
| BcM60C | MUC5AC_R2 | Tn | 188.0 (± 20.0) | 0.2 (± 0.0) | 1.1 (± 0.1) |
| | | C1 | 208.8 (± 36.0) | 0.1 (± 0.0) | 0.5 (± 0.1) |
| BcM60F | MUC5AC_R2 | Tn | 1.1 (± 0.2) | ND | ND |
| | | C1 | 1.0 (± 0.1) | ND | ND |
| | | 3SC1 | 1.4 (± 0.1) | ND | ND |
| BcM60M | MUC5AC_R2 | Tn | 413.9 (± 27.7) | 2.8 (± 0.5) | 6.7 (± 0.3) |
| | | C1 | 888.7 (± 52.7) | 14.4 (± 9.9) | 16.3 (± 2.0) |
| | | 3SC1 | 262.7 (± 9.4) | 5.6 (± 1.8) | 21.2 (± 1.3) |
| | | 6SC1 | 15.2 (± 4.7) | 0.05 (± 0.01) | 3.2 (± 0.4) |
| | | C2 | 610.9 (± 50.92) | 8.0 (± 5.6) | 13.1 (± 1.3) |

Errors are ± one-half of the 95% confidence interval.
<sup>1</sup> Determined using a simplified kinetic model. <sup>2</sup> ND, not determined.

The kinetic analysis of BcM60M gave k_cat_, K_M_, and k_cat_/K_M_ values of generally similar magnitude as for BcM60K and BcM60C **(Table 1)**. However, the order of glycan preference was different: C1 > C2 > Tn > 3SC1 > 6SC1. There was a 20-fold difference between the k_cat_/K_M_ for most and least preferred substrates with very low activity on the substrate having the 6SC1 glycan. There was roughly a two-fold preference for substrates with C1 compared with 3SC1, consistent with the FRET screen. We also generated a Y620F mutant of BcM60M, which showed a similar trend of preference: C1 > C2 ≈ Tn > 6SC1 **(Table 2)**. Except for the substrate having the 6SC1 glycan, on which the mutant had higher catalytic efficiency than the wild-type, the Y620 mutant had slightly lower overall activity relative to the wild-type.

**Table 2:** Kinetic parameters of the BcM60M Y620F mutant on the MUC5AC_R2 linker.

| O-glycan | $k_{\text{cat}}/K_M$<br>($\text{min}^{-1} \text{mM}^{-1}$) | Relative to<br>WT BcM60M |
| --- | --- | --- |
| Tn | 285.9 ($\pm 16.0$) | 0.7 |
| C1 | 610.9 ($\pm 50.9$) | 0.7 |
| 6SC1 | 20.4 ($\pm 1.6$) | 1.3 |
| C2 | 333.6 ($\pm 20.7$) | 0.5 |

Errors are ± one-half of the 95% confidence interval.

Also consistent with the FRET screen, BcM60F reactions progressed very slowly and we were only able to estimate k_cat_/K_M_ values by using the simplified hit-and-run analysis of the kinetic data. BcM60F did not display any obvious preference for Tn, C1, or 3SC1 **(Table 1)**.

### Quantitative analysis of BcM60K peptide selectivity

To better understand the influence of the peptide sequence on the efficiency of *O*-glycopeptidase cleavage we took the approach of generating hybrid linkers whereby the sequences before and after the glycosylation site in a “good” substrate (rapid cleavage) were swapped with the sequences before and after the glycosylation site in a “bad” substrate (no or slow cleavage). We chose the MUC5AC_R2 linker as the good substrate and the MUC4, MUC11, and IgA1 linkers as bad substrates (**Figure 5**), resulting in six Tn-modified hybrid linkers and three Tn-modified original linkers to compare (**Figure 5**). We used an initial assessment of BcM60K by FRET screen followed by detailed kinetic analysis (**Figure 5**). In the latter case the reactions were too slow for five of the substrates, thus the FRET screen and kinetic approaches proved complementary. The results consistently revealed that swapping in the N-terminal or C-terminal portion of the MUC5AC_R2 linker into the corresponding N- or C-terminus of a bad substrate improved cleavage of that substrate relative to the unaltered bad substrate. The N-terminal substitution has the largest effect where in the cases of MUC11 and IgA1, the improvement was quite profound going from inactivity to an efficiency roughly one-half and one-quarter, respectively, of the MUC5AC_R2 substrate. With MUC4, the hybrid substrates were roughly equally good substrates at about one-sixth that of the MUC5AC_R2 linker, whereas the enzyme was nearly inactive on MUC4.

**Figure 5.**
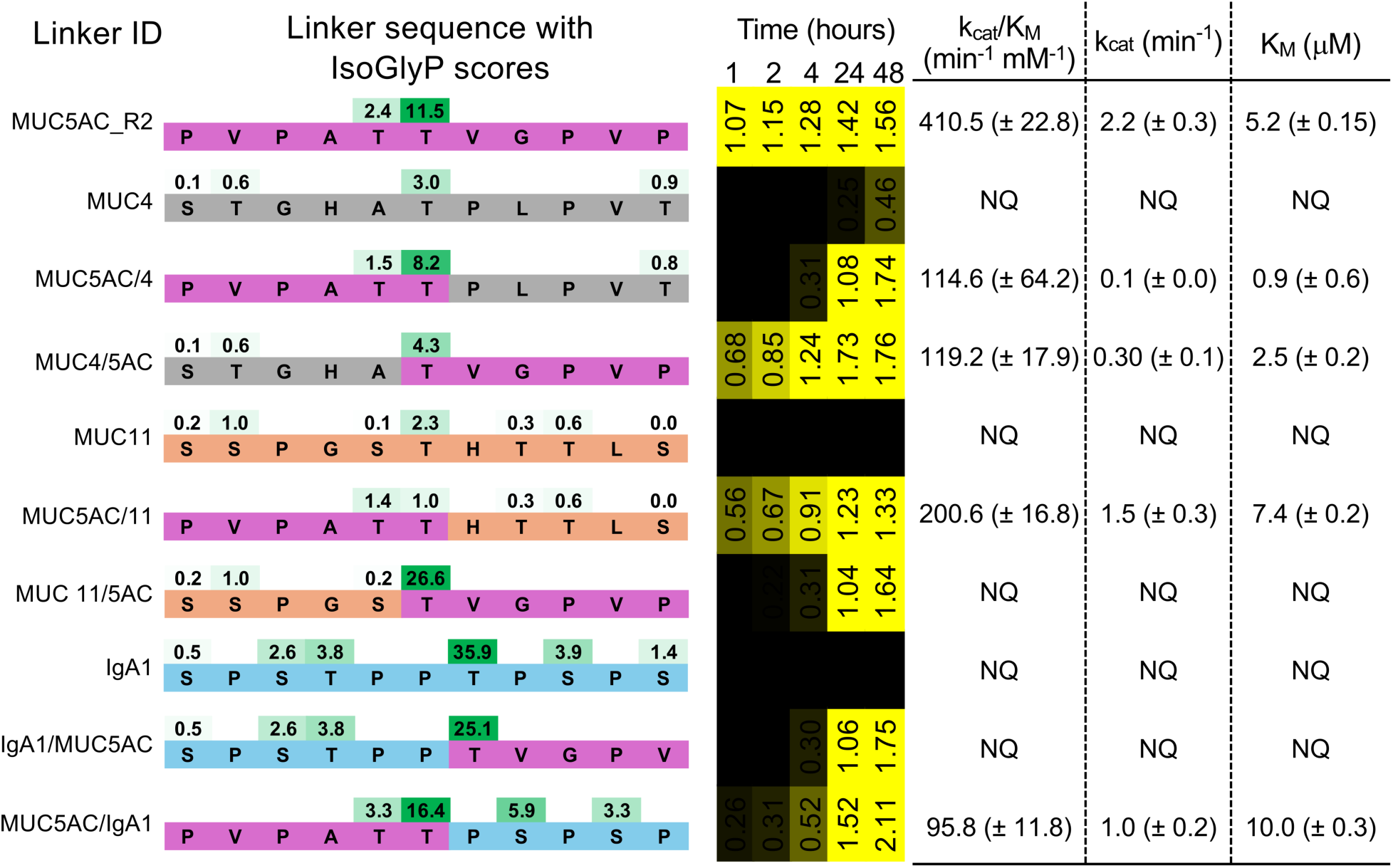
Analysis of BcM60K activity on hybrid substrates modified with the Tn antigen. The organization of the linkers in the FRET substrates are shown and color coded according to the origin of the sequence. IsoGlyP^41^ scores are provided to indicate the likelihood of a site being *O*-glycosylated by the GalNAcT2 isozyme; higher scores indicate higher likelihood. The middle column shows the FRET screen RoR for the substrates and the table the results of separate detailed kinetic studies.

### Structural analysis

To begin illuminating the structural features contributing to the specificity of the *B. caccae O*-glycopeptidases in clade 1, we determined the structures of BcM60B, BcM60F, BcM60K and BcM60C (**Figure 6**). The pairwise root mean square deviations (rmsd) roughly correlated with the pairwise amino acid sequence identities. The overall fold is that described for other M60-like *O*-glycopeptidases ^1320^, though BcM60B lacks the C-terminal β-sheet domain present in the other three enzymes. The active sites, identified by bound Zn^2+^ atoms, showed three different patterns of contouring in the predicted G-sites. BcM60C and K had very similarly shaped G-sites. BcM60B appeared to have an occluded G2’’ site whereas BcM60F had a slightly pinched G2’’ site and a more positive overall charge around the G2’ site.

**Figure 6.**
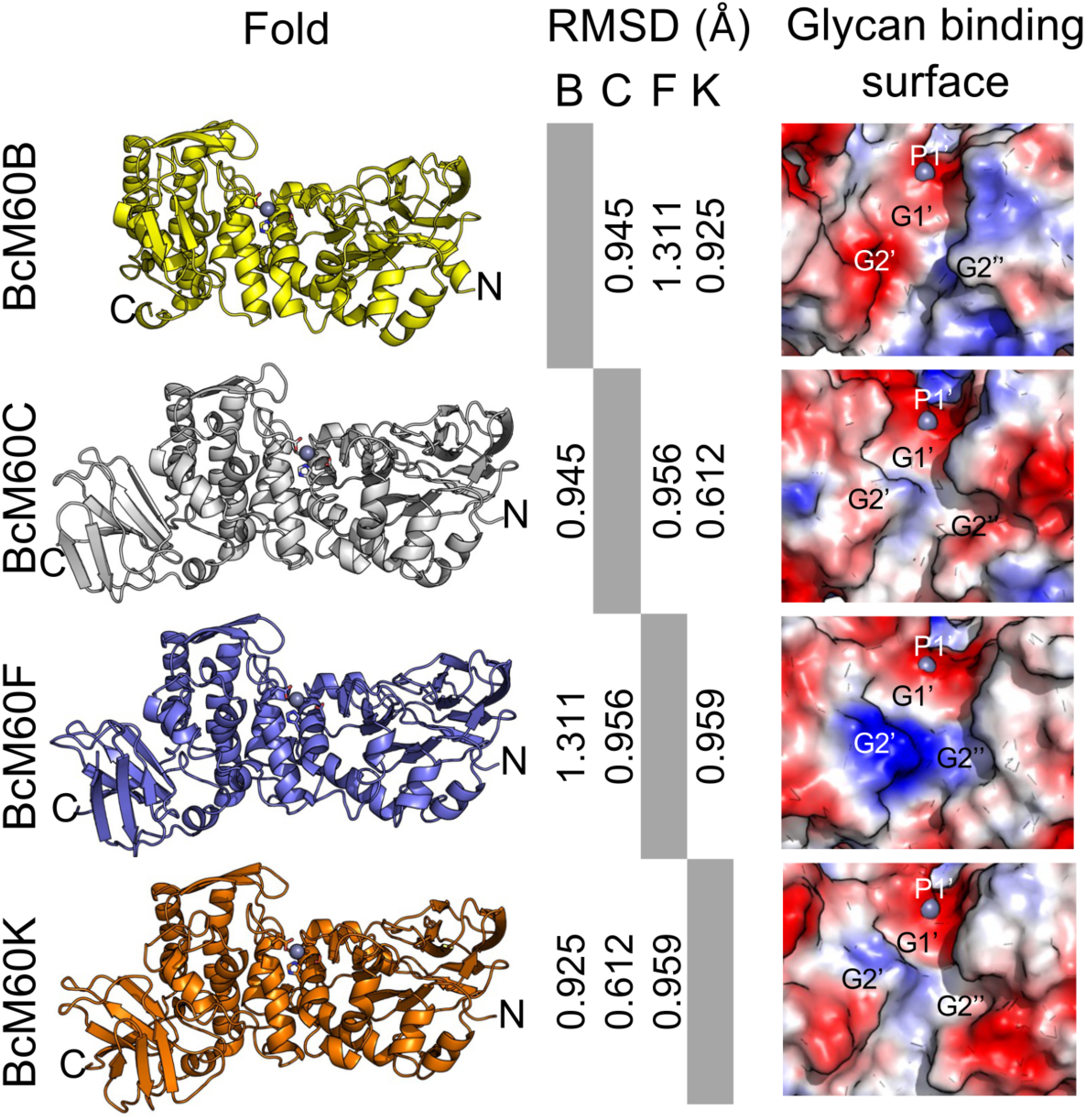
Structural analysis of BcM60s. The structures are shown in the context of a pairwise structural root mean square deviation matrix and surface representation of the active sites. The surface figures indicate a bound Zn^2+^ as grey sphere and putative active site subsites are labeled according to the nomenclature of Noach et al. ^13^.

Glyco-amino acids (GAA), such as 1-*O*-serinyl-GalNAc (GalNAc-Ser or Tn antigen), sometimes display the ability to bind to *O*-glycopeptidases if the enzyme accommodates it as a substrate fragment ^13^. We were unable to generate complexes of BcM60B and BcM60F, but we were able to determine the structures of both BcM60C and BcM60K in complex with C2-Thr and 6SC1-Ser. In all four cases the electron density was sufficiently defined to allow modelling of the entire ligands, though the electron density was sometimes incomplete for the GlcNAc or Neu5Ac occupying the G2’’ site, indicating some potential mobility (**Supplementary Figure 6**). The C2-Thr bound to both enzymes in essentially identical conformations with the same constellation of interactions (**Figure 7A**). The surface of BcM60C reveals the pocket forming the G2’’ site and how the β-1,3- linked galactose climbs up the sidewall of the active site channel (**Figure 7B**). Similarly, the 6SC1-Ser also bound to both enzymes in essentially identical manners (**Figure 7C**). In contrast to C2, however, the Neu5Ac pyranose ring in the G2’’ site was oriented at roughly right angles to the GlcNAc of C2 (**Figure 7D**). This orientation allows O9 of the Neu5Ac glycerol moiety to point down into a small pocket at the beginning of the catalytic helix, which contains the HEXXH motif. This results in hydrogen bonding between O9 and a backbone amide nitrogen and carbonyl oxygen (**Figure 7C**). In addition to shape complementarity of the glycan binding sites to the trisaccharide GAAs, all three monosaccharides in the glycan are involved in the hydrogen bonding network between the active site and the GAA.

**Figure 7.**
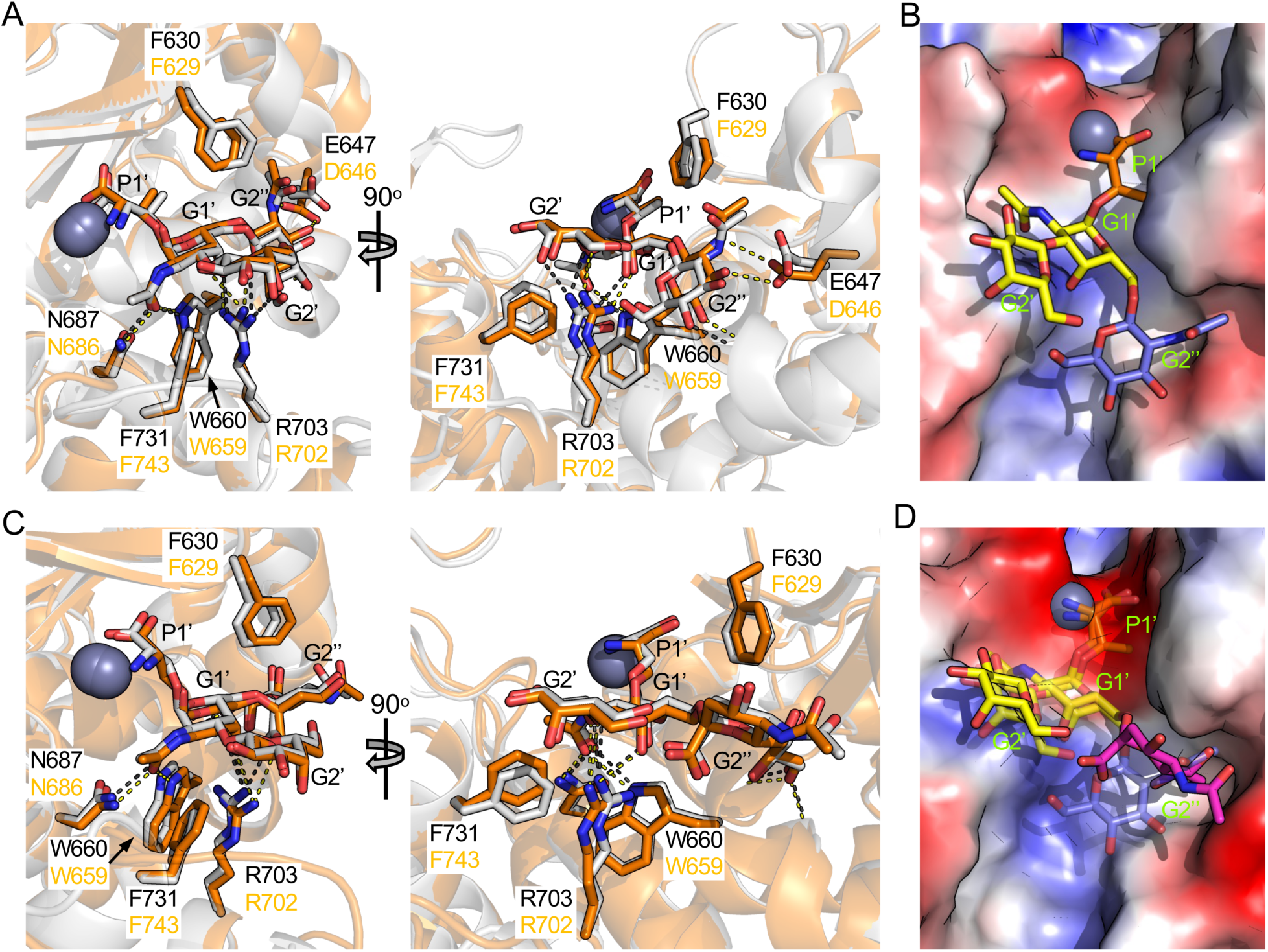
Structural analysis of BcM60C and BcM60K complexes. A) Structures of BcM60C (grey) and BcM60K (orange) in complex with C2-Thr. Potential hydrogen bonds are shown as dashes. Ligands are shown as sticks. B) The surface of the BcM60C active site colored according to charge (color ramped blue to white to red, positive to neutral to negative) showing the poise of the C2-Thr ligand and the G2’’ site. C) Structures of BcM60C (grey) and BcM60K (orange) in complex with 6SC1-Ser. Potential hydrogen bonds are shown as dashes. Ligands are shown as sticks. D) The surface of the BcM60K active site colored according to charge (color ramped blue to white to red, positive to neutral to negative) showing the poise of the 6SC1-Ser ligand and the G2’’ site. The C2- Thr ligand is shown as transparent sticks to show the different conformation of the GlcNAc and Neu5Ac groups in the G2’’ site.

We also co-crystallized a mutant of BcM60K that was catalytically inactivated by a E664A mutation with a MUC5AC glycopeptide (GTTPSPVPTTS**<u>T</u>**TSAP where the bold underlined residue bears an *O*-linked GalNAc). The electron density allowed modelling of the five P2 to P3’ amino acids and the GalNAc residue (**Supplementary Figure 6**). The P2 to P1’ residues of the substrate made anti-parallel β-strand-like pairing interactions with β-strand lining the enzyme active comprising residues M630 to T633 (**Figure 8A and 8B**). The P3’ residue packed against the phenyl ring of F629. The GalNAc residue made the same hydrogen bonding interactions observed for the core GalNAc in the C2-Thr and 6SC1-Ser complexes. The poise of the peptide is consistent with the catalytic mechanism proposed for clan MA metallopeptidases (**Figure 8A and 8B**)^21^. The carbonyl oxygen of the P1 residues interacts with the Zn^2+^ while a water molecule is positioned appropriately to attack the carbonyl carbon of P1. However, in the mutant, the catalytic glutamate (E664) is missing. The position of this residue was determined by overlapping a wild-type structure with the mutant complex. This revealed the catalytic glutamate to be in a position appropriate for activating the water molecule. The peptide binding region of the BcM60K active site has a trench-like shape, similar to that of AMUC_0627 (**Figure 8C**) ^22^. While many active site features are conserved between BcM60K and AMUC_0627, the latter also has the demonstrated presence of G1 and G2 glycan binding sites, which may involve non-conserved tryptophan (W149) and tyrosine (Y287) residues that line the peptide binding region of the active site (**Figure 8D**).

**Figure 8.**
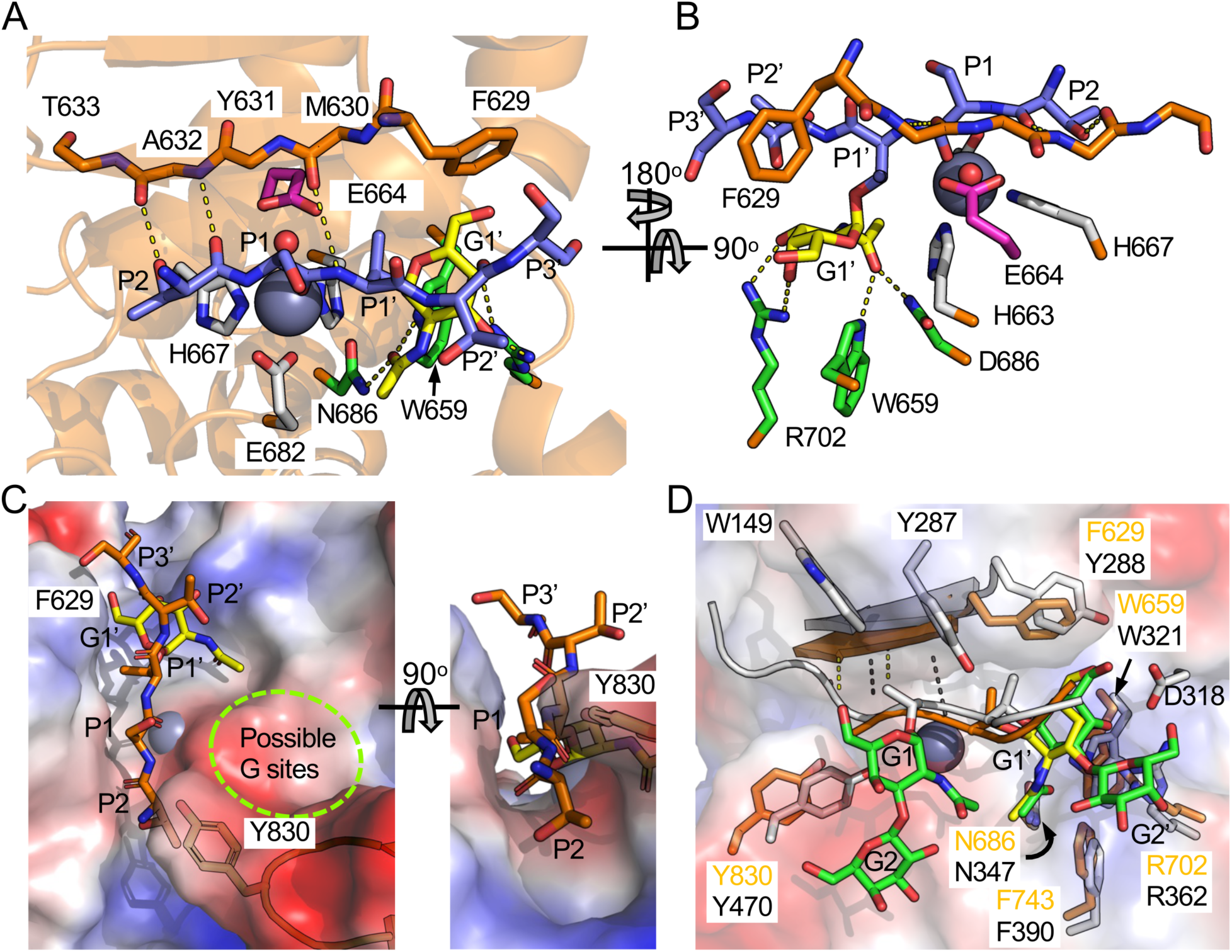
Structural analysis of BcM60K E664A mutant in complex with an intact MUC5AC glycopeptide. A) and B) Views of the active site with bound glycopeptide (orange and yellow sticks). The backbone of the β-strand lining the wall of the active site is shown as yellow strands. The zinc sites are shown in grey sticks, and sugar binding residues as green sticks. The catalytic glutamate is shown in pink and place by overlap of the wild-type structure with the mutant complex. Potential hydrogen bonds are shown as dashes. C) The surface of the BcM60K active site colored according to charge (color ramped blue to white to red, positive to neutral to negative) showing the poise of the glycopeptide ligand. D) Overlap of the BcM60K E664A glycopeptide (orange) complex with the AMUC_0627 E326A glycopeptide complex (grey; PDB ID 7YX8). The β-strand lining the active site is shown as a cartoon. Relevant side chains are shown as sticks.

## DISCUSSION

The five *B. caccae* peptidase_M60 *O*-glycopeptidases from clade 1 all display mucinase activity on bovine submaxillary mucin, which is related to human MUC19, but displayed differential selectivity in the FRET screen. Overall, BcM60C, BcM60K, and BcM60M accommodated linear and branched *O*-glycans and had similar selectivity for linker sequences. BcM60F recognized only linear glycans and a very limited set of linker sequences. Finally, BcM60B was inactive in the FRET screen and therefore is selective for a yet undefined glycan and/or peptide sequence.

All the enzymes examined, except for BcM60B, had activity on substrates with C1 glycans, thus indicating potential binding of the galactose residue in a G2’ site. However, none of the enzymes displayed significantly improved activity on C1 glycans relative to the Tn modified versions. Likewise, BcM60F also displayed no significant preference for the MUC5AC_R2 linker with 3SC1 compared with the C1 version, while BcM60M showed a two-fold decline in efficiency. BcM60K showed a more profound deleterious effect from the terminal sialic acid at around a 4-5 fold decrease in efficiency, while BcM60C was qualitatively poor on the sialylated substrates. An examination of the surface features of BcM60C, BcM60K, and BcM60M in the region of the predicted G3’ subsite indicates them to be acidic or neutral (**Figures 9A and 9B, and Supplementary Figure 7**), respectively. In contrast, the predicted G3’ subsite is quite basic in BcM60F (**Figures 9C**). A possible explanation for the reduced activity of BcM60K, and possibly BcM60C and BcM60M, on substrates bearing 3SC1 is the lack of charge complementarity between the G3’ subsite and the negatively charged Neu5Ac residue in the 3SC1 glycan (**Figures 9A and 9B**). Better charge complementarity is present in BcM60F, possibly leading to improved binding (**Figure 9C**).

**Figure 9.**
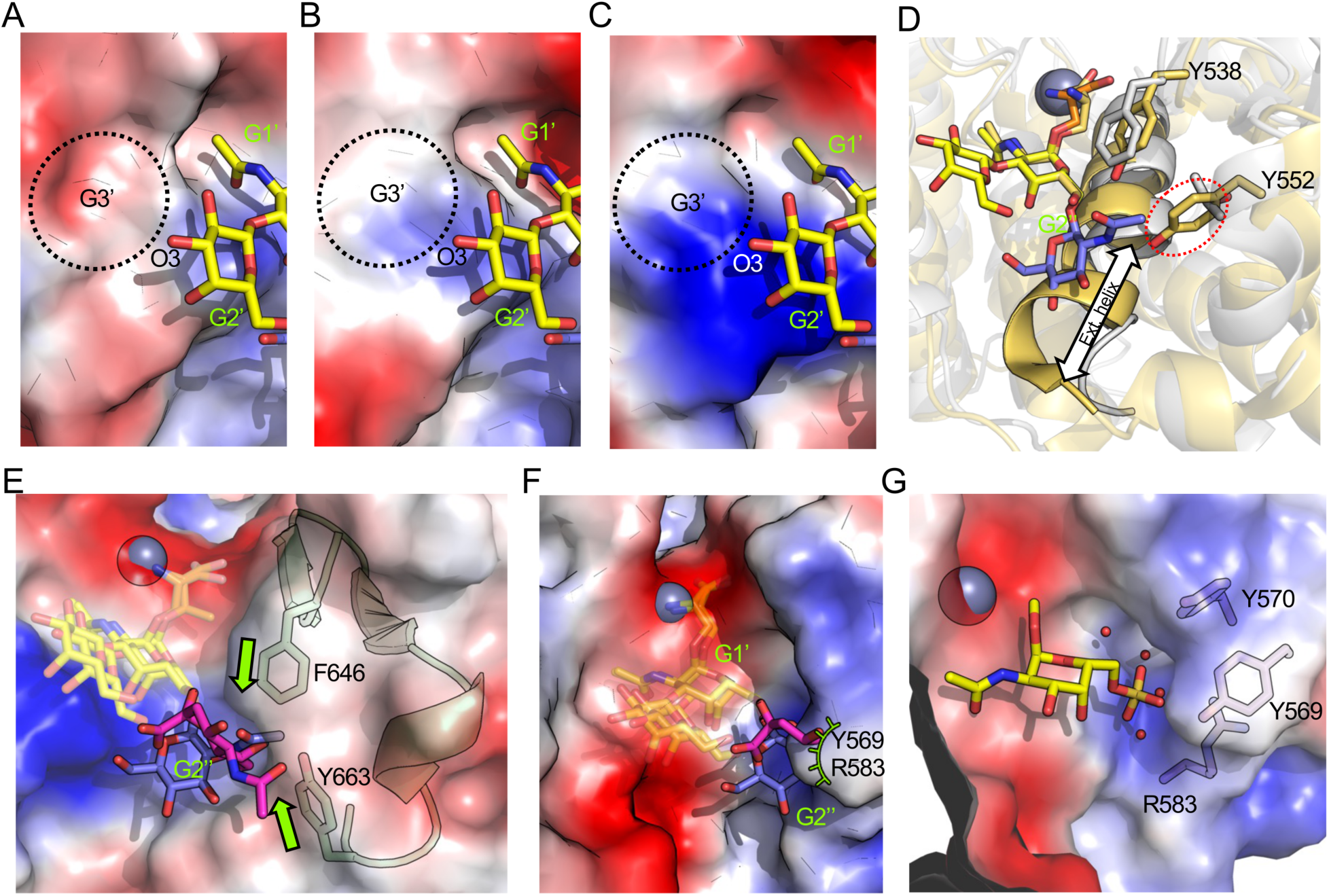
The structural basis of glycan recognition by the BcM60s. A-B) Surfaces of BcM60C, BcM60K, and BcM60F, respectively, colored according to charge (color ramped blue to white to red, positive to neutral to negative). The C2-Thr complexes of BcM60C and BcM60K are shown. The ligand was placed in BcM60F by an overlap with BcM60C. The regions where terminal α-2,3-linked Neu5Ac residues would lie are indicated as G3’ sites. D) Overlap of the BcM60C C2-Thr complex (grey) with the BT4244 Tn antigen complex (gold; PDB ID 5KD8). The arrow highlights the longer catalytic helix in BT4244 (see also **Supplementary Figure 8**). Relevant side chains in the G2’’ site are shown as sticks. E) Surface of the BcM60F active with the C2-Thr and 6SC1-Ser ligands placed by overlap with BcM60C. The shown secondary structure and sidechains of BcM60F highly pinching of the G2’’ site. F) Surface of the BcM60B active with the C2-Thr and 6SC1-Ser ligands placed by overlap with BcM60C. This highlights occlusion of the G2’’ site by residues Y569 and R583 in BcM60B. G) view of the BcM60B surface showing the basic water filled pocket formed by Y570, Y569, and R583. 1-O-methyl-6-sulfo-GalNAc was modeled in by first overlapping BcM60B with the BcM60C C2-Thr complex then aligning the sulfated GalNAc with the core GalNAc of the C2-Thr ligand.

It was previously postulated that a shortened helix containing the catalytic machinery correlates with the ability to accommodate branched glycans ^20^. The clade 1 *O*- glycopeptidases from *B. caccae* all have shortened catalytic helices relative to BT4244 and AMUC_0627 (**Figure 9D and Supplementary Figure 8**). In BT4244 (**Figure 9D**) and AMUC_0627 any potential G2’’ site is blocked by the lengthened helix and additional amino acid sidechains that fill the pocket. Though BcM60F has a shortened catalytic helix, the G2’’ pocket is pinched closed, largely due to the proximity of the F646 sidechain to Y663 (**Figure 9E**). The structures of BcM60C and BcM60K clearly show the G2’’ subsite pocket, the creation of which is partly enabled by shortened helix and the pocket being lined by short sidechains. An AlphaFold 3 generated model of BcM60M reveals a G2’’ subsite similar to that of BcM60K and BcM60C (**Supplementary Figure 7**). However, the clear preference that BcM60M had for C2 glycans over 6SC1 glycans (∼20-fold) distinguishes this enzyme from BcM60K, which only had an approximately 4-fold preference for C2 glycans. The most noticeable difference between the G2’’-sites of the two enzymes is that F629 in BcM60K is replaced by Y620 in BcM60M. The Y620F mutant of BcM60M did not have a profound difference in its preference for substrates with Tn or C1 glycans. However, while the wild-type enzyme had a 40-fold preference for substrates with C2 compared with 6SC1 glycans, the Y620F mutant only had a 17-fold preference for substrates with C2. This suggests that the presence of tyrosine *vs*. phenylalanine has some role in discriminating between C2 and 6SC1. The failure to restore the 4-fold selectivity of BcM60K, however, suggests that the differences in C2 and 6SC1 recognition by BcM60K and BcM60M is more nuanced than just the presence of tyrosine or phenylalanine in the G2’’-subsite.

BcM60B has appropriately placed metallopeptidase catalytic machinery as well as the key residues for *O*-GalNAc recognition. Though it displayed weak mucinase activity it was not active on any of the substrates in our FRET screen, making it an outlier. Though it has a short catalytic helix, the G2’’ position is completely occluded by Y569 and R583 and it lacks the conserved Phe/Tyr residue that is on the boundary of the G1’ and G2’’ sites of other M60 peptidases (*e.g.*, F629 in BcM60K; **Figure 9E**). These alterations, while blocking a potential G2’’ site, create a small positively charged water filled pocket adjacent to the G1’ site (**Figure 9G**). This pocket is reminiscent of sulfate binding sites in enzymes that are active on sulfated polysaccharides ^23^. To explore this, we generated an energy minimized 1-*O*-methyl 6-sulfo-GalNAc monosaccharide with GlyCam ^24^ and modelled it into the active site with guidance from an overlap of BcM60B with the BcM60C C2-Thr complex. By overlapping the 6-sulfo-GalNAc with the GalNAc of the C2 and with no alteration of the 6-sulfo-GalNAc at all, the sulfate group projected into the side-pocket of the BcM60B G1’ site with no clashes (**Figure 9G**). Based on this shape and charge complementarity we propose that BcM60B may recognize linear *O*-glycans with a 6-sulfo- GalNAc linked to the protein backbone. This modification is very rare, but has been identified on biologically relevant glycoproteins, such as salivary MUC7, and may be associated with the inflammatory state of the host ^25,26^. Selectivity for such a modified sugar would explain the very poor apparent mucinase activity of BcM60B and its lack of activity in our FRET screen.

The enzyme-to-enzyme differences that change the active site architecture to selectively bind linear glycans or also accommodate branched glycans is a complex interplay of multiple structural features that lead to nuanced substrate selectivity. The peptide portion of the substrate is another key aspect of substrate recognition, though peptidase_M60 *O*- glycopeptidases have been largely thought to be quite impartial to the amino acid sequence of their substrates. Only in the case of IMPa has a recent systematic semi-quantitative analysis shown the activity of this enzyme to be sensitive to the nature of the P1 amino acid ^27^. By using a broad range of well-defined substrates that are compared side-by-side, the studies here reveal that the peptide sequence of the substrate can profoundly influence enzyme activity. While some of the linker sequences in the FRET screen have multiple potential *O*-glycosylation sites, and thus glycosylation pattern may be a factor, several substrates have only a single plausible predicted glycosylation site (*i.e.*, MUC1_R1, MUC1_R2, MUC5AC_R1, MUC4, MUC8_R1, and Fetuin) while the mass spectrometry results showed the MUC5AC_R2 and IgA1_h linkers to only bear single glycans. Despite only having single glycans, BT4244 clearly favored the MUC5AC_R2 substrate over the IgA1_h substrate. BcM60C, BcM60K, and BcM60M showed highly variable activity on the single glycan substrates: they were highly active on MUC1_R1, MUC5AC_R1, MUC5AC_R2; had medium activity on Fetuin; and low or no activity on MUC1_R2, MUC4, MUC8_R1, and IgA1_h. Finally, As the most obvious example of peptide sequence selectivity, BcM60F only had activity on the MUC5AC_R1 and MUC5AC_R2 linkers and nothing else. This is a strong indication that the amino acid sequence context of the *O*-glycosylation site has a strong influence in determining how efficiently *O*-glycopeptidases hydrolyze substrate.

This was further supported by the analysis of engineered linkers whereby the hybrids incorporated features of good *vs*. bad substrates. In all cases, swapping out the N- or C-terminal half of substrate pairs improved the activity of BcM60K on the hybrid relative to the bad reference substrate. In two out of the three cases, using an N-terminal half from the good substrate had the most profound effect on improving activity while in the third case (MUC5AC_R2 and MUC4 hybrids) the influence of swapping out the N- or C-terminal halves was roughly the same. Taken together, this indicates that amino acid sequences on either side of the P1’ residue (the P and P’ sides) play an important role in substrate recognition with the P side of the sequence perhaps having the most impact.

The structure of BcM60K E664A in complex with a MUC5AC derived Tn-glycopeptide revealed the β-strand pairing interaction between the peptide portion of the substrate and the enzyme active site, the trench-like architecture of the active site, and interactions involving both the P and P’ sides of the peptide sequence (**Figures 8A to 8C**). One wall of the active site trench is created by the β-strand that pairs with peptide portion of the substrate. The other wall of the trench is created by the encroachment of a loop containing Y830, which partially closes over the active site. The narrow active site leads to the question of how the peptide gains access to the catalytic machinery. We postulate this is enabled by protein motions that flex the active site enough to enable substrate access. Though the loop containing Y830 is in same position in the unliganded and peptide bound structures, the B-factors of the loop and the structures supporting the loop in the unliganded structure are significantly higher than those of the peptide bound structure (**Supplementary Figure 9**). Similarly, in BcM60C the equivalent loop to that containing Y830 in BcM60K has Y817 in the corresponding position. In the unliganded structure of BcM60C this loop could modeled in two partially occupied positions and in neither case could the tyrosine sidechain be modeled (**Supplementary Figure 10**). In one conformation, the tip of the loop is pulled out of the active site, substantially opening it up. Thus, we propose that active site malleability in these enzymes is necessary to allow the peptide backbone of the substrate to slot into the trench.

In the active site, the poise of the β-strand of the peptide substrate places the side chains in an alternating up and down orientation with the P1’ residue, which is necessarily a serine or threonine, in the down orientation (**Figure 8C**). The P1 residue is in the up conformation pointing into solvent, which suggests it could be an amino acid with a larger side chain or bear a P1 glycan that interacts with possible G sites. This is indeed the case with AMUC_0627 where Y470, which is conserved with Y830 in BcM60K, makes the base of the G1 site ^22^ (**Figure 8D**). Unlike AMUC_0627, however, BcM60K lacks the equivalents of W149 and Y287 of AMUC_0627, which seems to create an aromatic side wall that may be key to binding P1 glycans ^20,22^ (**Figure 8D**). The P2 residue is in the down position with the sidechain occupying a surface pocket. The limited dimension of this pocket suggests that very large sidechains, such as Trp, Tyr, Lys, and Arg would not fit in the S2 subsite (**Figure 8C**). Only one potential substrate, MUC7_R1, has the potential for a Lys in this position but it is unknown if the nearby potential site of glycosylation is or is not glycosylated or cleaved (**Supplementary Figure 11**). However, MUC6B_R1 has a Tyr in this position and was one of the substrates that was not cleaved (**Supplementary Figure 11**). The P2’ sidechain was not directly involved in any interactions and was exposed to solvent, suggesting potential tolerance of a wide variety of sidechains in this position. Finally, the backbone of P3’ packed against F629 and suggested that some residues may make more optimal interactions than others, but precisely how is unclear at this time.

With the observations that the substrate amino acid sequence plays a role in the efficiency of cleavage and the ensuing concept that the active site architecture should influence specific amino acid sequence recognition, we compiled and merged the FRET screen and kinetic date for BcM60K, BcM60C, and BcM60M to examine any potential patterns in substrate sequence and cleavage efficiency, which we scored as NA (no cleavage), poor, medium, and good (**Supplementary Figure 11**). Other than the one substrate, MUC6_R1, with a Tyr in the P2 position, we saw no obvious patterns. A weak pattern, however, is that the poor or uncleavable substrates are likely to have a proline at P1 or P2’ (or both) whereas the medium or good substrates are more likely to have a potential cleavage site with an absence of neighboring prolines. It appears from this analysis, that there are few cases where amino acids in a certain position are not tolerated, excepting P1’ of course.

The results herein continue to highlight the sophisticated nature of *O*-glycopeptidases. The combinatorial effect of peptide backbone features and glycan characteristics create the potential for extraordinarily diverse substrates. The structural features of individual peptidase_M60 enzymes appear to complement subsets of substrates thereby defining the unique selectivity of even closely related peptidase_M60 proteins. Indeed, we have shown that the five *B. caccae O*-glycopeptidases from clade 1 represent at least three different selectivities. These are five of the sixteen that *B. caccae* is known to deploy and we have shown an additional eight have mucinase activity. The remaining three *B. caccae* peptidase_M60 proteins are not characterized at any level but are presumably also functional. If one applies the concept shown here that not all enzymes from the same clade display identical selectivities, the arsenal of *B. caccae O*-glycopeptidases is likely to have a very broad diversity of substrates that it may target. The implications are that *B. caccae* is a very accomplished glycoprotein/mucin degrader and that this is enhanced by multiple *O*-glycopeptidases that target different substrate features. In other words, a single *O*-glycopeptidase alone may not have broad enough selectivity to do the job well. We anticipate that expansion of *O*-glycopeptidase encoding genes will become a marker for proficient mucin degrading bacteria, such as, for example, the known mucin degrader

*A. muciniphila*, which as two known M11-like and three known peptidase_M60 *O*- glycopeptidases ^14,20,22,28,29^. Overall, it highlights the potential diverse selectivities of even closely related peptidase_M60 *O*-glycopeptidases, which might otherwise look like redundant enzymes, and points to the potential need of accomplished mucin degrading bacterium to deploy numerous similar enzymes with different selectivities to achieve efficient mucin breakdown.

## EXPERIMENTAL PROCEDURES

### Materials

All reagents, chemicals and other carbohydrates were purchased from Sigma unless otherwise specified. 6SC1-Ser was purchased from Biosynth and the MUC5AC glycopeptide from Sussex Research Chemicals.

### Enzyme Purification and Production

Peptidase*_*M60 metallopeptidase modules were synthesized by GenScript or Twist Bioscience and codon optimized for expression in *Escherichia coli* (see **Supplementary Table 2** for protein names and locus tags). These genes fragments were cloned into pET28a by GenScript or Twist Bioscience. Competent *E. coli* BL21 (DE3) cells (Invitrogen) and Shuffle T7 cells (New England BioLabs) were transformed with the appropriate pET28a recombinant expression plasmid by heat shock. Transformed cells were spread plated onto LB agar plates with 50 µg/mL kanamycin to select for cells transformed with the pET28a plasmid. Plates were incubated overnight at 37°C. Selected colonies were used to inoculate 2xYT media supplemented with 50 µg/mL kanamycin and grown at 37°C until an optical density of ∼ 0.8 at 600 nm was reached. Cultures were cooled at 16°C for 1 hour and recombinant protein expression was induced with isopropyl-b-D-1-thiogalactopyranoside (IPTG) at a final concentration of 0.5 mM. Incubation was continued at 16°C with shaking overnight. Cultures were centrifuged at 6300 rpm for 10 min at 10°C in a Beckman Coulter Avanti J-E centrifuge. Supernatant was discarded and the cell pellet was disrupted by chemical lysis with the stepwise addition of sucrose solution (25% sucrose and 20 mM Tris-HCl pH 8), 0.66 mg/mL lysozyme, deoxycholate solution (1% deoxycholate, 1% TritonX100, 50 mM Tris-HCl pH8, 100mM NaCl), MgCl_2_ and DNase as previously described ^13^. Cell lysate was centrifuged at 16,500 rpm for 30 min at 7°C prior to purification.

Cleared cell lysate was loaded onto Ni^2+^-NTA immobilized metal affinity resin (IMAC) equilibrated with binding buffer (20 mM Tris-HCl pH 8, 500 mM NaCl, 10% glycerol). Nonspecific binding to the resin was eliminated with a 20 mM imidazole and His-tagged protein was eluted with 500 mM imidazole. Samples of each fraction were combined in a 1:1 ratio with 2X loading buffer and run on a 12% SDS-PAGE gel stained with Coomassie. Fractions containing pure protein were concentrated using an Amicon stirred ultrafiltration unit with a 10 kDa molecular weight cut-off (MWCO) membrane (Millipore). Proteins for activity assays were dialyzed overnight in binding buffer (20 mM Tris-HCl pH 8, 500 mM NaCl, 10% glycerol) at 10°C.

For the purpose of crystallization, the N-terminal six-histidine tag was cleaved from the proteins by restriction grade thrombin with 150 mM NaCl and 2.5 mM CaCl_2_ according to the manufacture procedure. Size exclusion chromatography using a Sephacryl S200 HR column (GE Healthcare) in 20 mM Tris-HCl pH8 and 100 mM NaCl was used to purify proteins lacking the N-terminal six-histidine tag. Fractions containing purified protein were pooled and concentrated using an Amicon stirred ultrafiltration unit with a 10 kDa MWCO membrane (Millipore). Proteins were further concentrated using a Centricon. Protein concentrations were determined with the absorbance at 280 nm, the molecular mass and the predicted extinction coefficient (**Supplementary Table 2**). NanH was produced and purified as described previously ^19^.

### Generation of FRET substrates

The initial template for our FRET substrates was ordered as synthetic gene with codon optimization for *E. coli* from Twist Biosciences. This comprised, in order from the N-terminus, a 6-his tag for purification, the mNeonGreen fluorescent protein domain, an eleven amino acid linker, and C-terminal mScarlet I domain. Twenty-four synthetic gene fragments comprising a portion of the mNeonGreen gene, the sequence of the desired linker, and a portion of the mScarlet I domain were ordered from IDT (see **Supplementary Table 1**). Infusion cloning was used to replace the linker sequence of the FRET template using standard procedures. Sequence fidelity was confirmed by whole plasmid sequencing (Plasmidsaurus). The same approach was used to generate the hybrid linkers.

Unglycosylated FRET substrates were produced in *E. coli* BL21 using the procedures described above. To produce Tn-modified substrates, competent *E. coli* Origami2 DE3 cells (Novagen) were double-transformed with the desired FRET construct plasmid and the pOGO 42 plasmid ^18^. Culture growth followed procedures as described except that media contained 50 μg/mL kanamycin and 35 μg/mL chloramphenicol. Cells were pelleted by centrifugation at 5500 rpm for 15 minutes at 4°C in a Beckman Coulter Avanti J-E centrifuge. Supernatant was discarded and cell pellets were resuspended in 20 mM HEPES pH 7, 10 mM MnCl_2_ and stirred for 40 minutes with 10 mg lysozyme, BugBuster® 10X stock to 1X final concentration, 0.2 mg DNAse I and MgCl_2_ to a final concentration of 5 mM. Cell lysates were centrifuged at 15500 rpm for 45 minutes to separate supernatant from cell debris. Supernatants were transferred to sterile tubes and sodium azide was added to 0.05% w/v to prevent bacterial grow-back. Uridine diphosphate *N-* acetylglucosamine (UDP-GlcNAc) was added to a final concentration of 15 mM. The reactions were covered in aluminum foil and incubated overnight at room temperature with rocking. Following overnight incubation, Tris-HCl pH 8 was added to a final concentration of 50 mM. The reactions were centrifuged at 10000 rpm for 10 minutes to pellet any grow-back, and cleared supernatant was used for purification.

Supernatant was loaded onto Ni^2+^-NTA IMAC resin (Thermo Scientific HisPur beads) equilibrated with binding buffer. The column was washed with three column volumes of binding buffer followed by two column volumes of 20 mM imidazole in binding buffer and 2 mL of 500 mM imidazole in binding buffer for elution of the His-tagged recombinant proteins. Elution fractions were concentrated on a stirred ultrafiltration unit (Amicon) using a 10 kDa molecular weight cut off membrane (EMD Millipore) and buffer exchanged Into 20 mM Tris pH 8 on Sephadex G-25 desalting columns (Cytiva). Installation of the Tn antigen was confirmed using lectin blotting.

The Tn-modified FRET substrates were sequentially converted by *in vitro* enzymatic reactions to C1, 3SC1, 6SC1, or C2 using approaches described previously ^13,28,30–32^.

Fmoc-Thr(β-D-Gal(1-3)[β-D-GlcNAc(1-6)]-α-D-GalNAc)-OH (Peracetate) was obtained from Sussex Research (Cat. #GA131030). Deprotection was performed under Zemplén conditions with 3.5 eq sodium methoxide; complete deprotection was confirmed by proton NMR. The resulting C2-Thr was purified by size exclusion in 0.1% aqueous formic acid to remove excess sodium then lyophilized.

### Substrate glycosylation analysis

All substrates used for lectin blotting were diluted to a concentration of 1 μM with 50 mM HEPES pH 7. 1 μL of each FRET substrate was spotted onto a 5 x 5 cm square of 0.45 μM nitrocellulose membrane (Bio-Rad Trans-Blot Transfer Medium). Confirmed substrates with the same glycosylation status were used as positive controls and unglycosylated substrates were used as negative controls. The spots were left until visibly dry. The membrane was then blocked for 30 minutes with rocking using a solution of PBS pH 7, 2% w/v bovine serum albumin, and 0.05% v/v Tween-20. The membrane was then washed three times with PBS + 0.05% Tween 20. The membrane was then incubated with rocking for 30 minutes with a PBS solution containing 10 μg/mL biotinylated lectin and 0.1 mM CaCl_2_, then washed twice with PBS. A solution of PBS + 0.05% Tween 20 and a final concentration of 5 μg/mL of Streptavidin Alexa Fluor^TM^ 790 conjugate (Invitrogen) was added to the membrane and incubated in a tin foil wrap with rocking for 30 minutes. Finally, the membrane was washed three times with PBS + 0.05% Tween 20, and two times with PBS. Imaging of membrane was done on the Mandel Li-Cor Odyssey CLx imaging system at a wavelength 800 nm. Lectin blots used *Vicia villosa* lectin (VVA) for Tn antigen, peanut agglutinin (PNA) for Core 1 glycosylation, and a lack of PNA signal for 3SC1 modified substrates. Biotinylated VVA and PNA were purchased from Vector Laboratories.

For molecular weight determination reverse phase high performance liquid chromatography followed by detection using mass spectrometry (RP-HPLC-MS) was performed using an Agilent 1200 SL HPLC System with a Phenomenex Aeris 3.6um, WIDEPORE XB-C8, 200Å, 2.1x50mm with guard column. An aliquot of the sample was injected onto the column at a flow rate of 0.45mLmin^-1^ and 95% mobile phase A (0.1% formic acid in water) and 5% mobile phase B (0.1% FA in acetonitrile). Elution of the analytes was done by using a linear gradient from 5% to 10% mobile phase B within 1 minute, 10% to 65% mobile phase B over a period of 6.5 minutes, 65% to 98% mobile phase B over a period of 0.5 minute, kept at 98% mobile phase B over a period of 1 minute and back to 5% mobile phase B over a period of 1 minute. Column temperature was 40°C. Mass spectra were acquired in positive mode of ionization using an Agilent 6220 Accurate-Mass TOF HPLC/MS system (Santa Clara, CA, USA) equipped with a dual sprayer electrospray ionization source with the second sprayer providing a reference mass solution. Mass correction was performed for every individual spectrum using peaks at m/z 121.0509 and 922.0098 from the reference solution. Mass spectrometric conditions were drying gas 10 L/min at 325°C, nebulizer 28 psi, mass range 100-3200 Da, acquisition rate of ∼1.07 spectra/sec, fragmentor 215V, skimmer 65V, capillary 4000V, instrument state 4GHz High Resolution. Data analysis was performed using the Agilent MassHunter Qualitative Analysis software package version B.03.01 SP3.

### Mucinase assay

The mucinase assay was performed as described previously ^13^. Briefly, microplate wells (96 well, MICROLON, high binding, Greiner bio-one) were coated with biotinylated bovine submaxillary mucin (0.08 µg/mL in PBS, 100 µL/well) and incubated overnight at 4°C. Wells were washed with PBS (3 x 100 µL) and incubated with 2 µM protease in PBS, 1% BSA and 0.5 mM ZnCl_2_ (100 µL/well) overnight at 37°C. Proteases were removed and wells were washed with PBS (4 x 200 µL). Free binding sites were blocked with 1% BSA in PBS with 0.1% Tween-20 (PBST, 200 µL/well) overnight at 4°C. Wells were washed with PBST (2 x 200 µL) and biotinylated substrates were detected with streptavidin-peroxidase solution (1:20000 dilution) freshly prepared in PBST. Wells were incubated with the streptavidin-peroxidase solution (100 µL/well) for 1 hour at room temperature and washed with PBS (4 x 200 µL). *O*-phenylenediamine dichloride (100 µL/well) was added as the peroxidase substrate. Upon colour appearance, the reaction was stopped by 0.5 M sulfuric acid and absorbance was measured at 492 nm with a Spectramax M5 plate reader using SoftMax Pro-6.2.1 software. Each reaction was tested in triplicate.

### FRET activity screens

FRET activity assays were performed essentially as described previously ^18^. Activity screens were setup in 384 well low-profile, clear bottom, black side microplates. 5 μL of each substrate at 0.4 mg/mL (7.2 μM) in reaction buffer (50 mM HEPES, pH 7.0) was dispensed into the tray. Reactions were initiated by the addition of 5 μL of enzyme at double the desired final concentration. Enzyme stocks were in reaction buffer with 200 μM ZnCl_2_. Samples reactions were prepared in duplicate. A paired set of controls were included and contained substrate but only reaction buffer with 200 μM ZnCl_2_ was added. Reads were taken at room temperature (22 °C) by excitation at a wavelength of 430 nm with measurement of emission intensity at 518 nm (F_518_) and 590 nm (F_590_) on a SpectraMax M5. Reads were at 1, 2, 4, 24, 48, and 72 hours. Data were processed by first averaging the duplicate reads then calculating the ratio of F_518_/F_590_ (F_R_) for samples and matched controls. A change in this ratio indicates a change in FRET, which was better revealed by taking the ratio of the F_R_ values and subtracting 1, to give ΔFRET (ΔFRET = F_R,sample_/F_R,control_ – 1). We refer to this as the Ratio-of-Ratios or RoR approach. An increase in ΔFRET indicates a decrease in FRET due to cleavage of the substrate. This was displayed as a heat-map vs. time with a ΔFRET threshold of 1 (*i.e.*, a doubling of the signal) assigned yellow and values below 0.2 colored black.

Substrate-enzyme combinations showing a significant ΔFRET (>0.4 at 24 hours) were flagged for further analysis. In this case, using the averaged raw fluorescence intensities, a fluorescence difference was calculated as F_518_ – F_590_. A change in fluorescence normalized to the controls, ΔF, was calculated as (F_518_-F_590_)_sample_-(F_518_-F_590_)_control_. ΔF was plotted vs time, and the progress curves analyzed using DynaFit with a method that has been referred to as “hit-and-run” ^33^. This reduces the reaction to a simple bimolecular reaction (E + S → E + P) with a single rate constant that approximates k_cat_/K_M_. The molar response for ΔF was allowed to float, eliminating the need to determine this independently, with the assumption that 100% of the substrate had the potential to cleaved. A y-offset was also included as a fit variable to account for slight differences in base fluorescence of the controls *vs*. the samples.

Studies of the synergy between BcM60K and NanH on 3SC1 substrates were performed and analyzed as above except with plus/minus the addition of 50 nM NanH at the time of initiating the reactions.

### Enzyme kinetics

Individual reactions were setup identically to the approach used for screening in a 10 μL final volume. We used a range of eleven final substrate concentrations from 0.25 – 6 μM. Reactions were setup using 5 μL of substrates containing 200 μM ZnCl and initiated by the addition of 5 μL of 2x enzyme stocks at concentrations of 0.02 μM, 0.2 μM, 0.4 μM, 0.5 μM, and 1 μM for BT4244, BcM60K, BcM60C, BcM60M, and BcM60F, respectively. A set of matched controls with no enzyme was included. All reaction and control samples were setup in duplicate. Reads were taken with the same parameters as for the screen except for the read interval (∼60 seconds) and the total kinetic run time (∼4 hours, or ∼18 hours for BcM60F). The raw fluorescence intensities of the controls were averaged but the duplicate reaction measurements were kept separated. A reaction progress of ΔF vs time for each set of reaction measurements were calculated as (F_518_-F_590_)_sample_-(F_518_-F_590_)_control_. This resulted in each enzyme concentration giving two sets of progress data comprising eleven reactions, for a total of 22 progress curves. Using DynaFit, this data was fit with the Van Slyke-Cullen model ^3334^. The same parameters used for fitting the screen data were used for this analysis with the addition of both k_cat_ and k_cat_/K_M_ as fit parameters. We were unable to do full fits for the BcM60F kinetic data as k_cat_ gave nonsensical values; therefore, we used the hit-and-run approach to estimate the k_cat_/K_M_. The data sets for each enzyme were analyzed using global fits of all the progress curves.

### Crystallization, diffraction data collection and processing

Crystallization conditions for all proteins were initially screened using up to 10 different sparse matrix screens and the sitting drop method of vapor diffusion in 96-well plates. Optimizations were carried at 18°C by hanging drop vapor diffusion. Relevant protein parameters and crystallization conditions are given in **Supplementary Table 3**. Complexes of BcM60C and BcM60K were obtained by soaking crystals in crystallization solution containing 20 mM ligand. Note that the BcM60C E665Q mutant was used to obtain the C2-Thr complex. BcM60K E664A was co-crystallized with 20 mM glycopeptide in the condition given in **Supplementary Table 3**.

Diffraction data were collected on an instrument comprising a Pilatus 200K 2D detector coupled to a MicroMax-007HF X-ray generator with a VariMaxTM-HF ArcSec Confocal Optical System and an Oxford Cryostream 800. Data were integrated, scaled and merged using HKL2000 ^35^. Data processing statistics are shown in **Supplementary Table 4**.

### Structure solution and refinement

All structures were determined by molecular replacement using AlphaFold 3 models as search models in PHASER. Refinements were carried out by a combination of REFMAC and Phenix.refine ^36,37^. The initial model was manually corrected using COOT followed by refinement in an iterative manner ^38^. For all structures, the addition of water molecules was performed in COOT with FINDWATERS and manually checked after refinement. In all datasets, refinement procedures were monitored by flagging 5% of all observations as “free” ^39^. Model validation was performed with MOLPROBITY ^40^. Model refinement statistics are shown in **Supplementary Table 4**.

## ACKNOWLEDGEMENTS

This research was supported by a Canada Institutes for Health Research Project Grant (FRN 04355). Additional financial support was from GlycoNet, NMR and mass spectrum data was collected by GlycoNet Integrated Services at the University of Alberta.

## AUTHOR CONTRIBUTIONS

BP, KB, AD, and ABB performed crystallography experiments; BP and AD performed mucinase assay; KB, OC, LM, and NT generated glycosylated FRET substrates; KB, BA, LM and OC performed FRET activity screens and kinetic analyses; KB performed sialidase synergy experiments; ABB conceived experiments and wrote draft manuscript; WW and ABB obtained funding, supervised, and edited manuscript.

## ACCESSION CODES

The atomic coordinates for the crystal structures reported here have been deposited in the Protein Databank under the accession codes 9PKS, 9PKT, 9PLX, 9PLW, 9PL7, 9PM5, 9PL8, 9PLA, and 9PM4.

