## Supplementary material for "Substrate recognition and cleavage by mucin degrading *O*-glycopeptidases from the gut microbe *Bacteroides caccae*"

Running title: *B. caccae O*-glycopeptidases

*To whom correspondence should be addressed: Alisdair B. Boraston, Department of Biochemistry and Microbiology, University of Victoria, Victoria, British Columbia, Canada, V8P 5C2;; Tel. +1 (250) 472-4168; Fax. +1 (250) 721-8855

**Supplementary Table 1.** Linker sequences in FRET substrates and DNA fragments used to clone them. Green nucleotide indicates the 3’ region of the mNeonGreen domain, red the 5’ region of mScarletRed 1, and black the linker coding sequence.

| Linker | Sequence | DNA Fragment |
|---|---|---|
| MUC1_R1 | PAPGSTAPPAH | CCGATGGCCGCAAACTACCTTAAGAACCAGCCCATGTACGTGTTTCGGAAAACTGAGCTTAAACACAGCAAGACAGAGTTAAATTTCAAGGAGTGGCAAAAGGC<br>ATTCACCGACGTTAAGCCTGCACCGGGAAGCACGGCTCCCCCTGCTCATGTGAGCAAAGGTGAGGCCGTGATCAAGGAATTCATGCGTTTCAAGGTCC<br>ACATGGAAGGATCAATGAACGGACACGAATTCGAGATAGAAGGGGAAGGCGAAGGTCGTCCATACGAAGGTACGCAAACAGCAAAACTGAAGGTGACG |
| MUC1_R2 | AHGVTSAPDTR | CCGATGGCCGCAAACTACCTTAAGAACCAGCCCATGTACGTGTTTCGGAAAACTGAGCTTAAACACAGCAAGACAGAGTTAAATTTCAAGGAGTGGCAAAAGGC<br>ATTCACCGACGTTAAGGCGCATGGGGTGACCTCAGCCCCTGACACACGTGTGAGCAAAGGTGAGGCCGTGATCAAGGAATTCATGCGTTTCAAGGTCC<br>ACATGGAAGGATCAATGAACGGACACGAATTCGAGATAGAAGGGGAAGGCGAAGGTCGTCCATACGAAGGTACGCAAACAGCAAAACTGAAGGTGACG |
| MUC2_R1 | TTVTPTPTPTG | CCGATGGCCGCAAACTACCTTAAGAACCAGCCCATGTACGTGTTTCGGAAAACTGAGCTTAAACACAGCAAGACAGAGTTAAATTTCAAGGAGTGGCAAAAGGC<br>ATTCACCGACGTTAAGACCACGGTTACGCCTACCCCGACGCCTACTGGCGTGAGCAAAGGTGAGGCCGTGATCAAGGAATTCATGCGTTTCAAGGTCC<br>ACATGGAAGGATCAATGAACGGACACGAATTCGAGATAGAAGGGGAAGGCGAAGGTCGTCCATACGAAGGTACGCAAACAGCAAAACTGAAGGTGACG |
| MUC2_R2 | VPPTTTPSPPP | CCGATGGCCGCAAACTACCTTAAGAACCAGCCCATGTACGTGTTTCGGAAAACTGAGCTTAAACACAGCAAGACAGAGTTAAATTTCAAGGAGTGGCAAAAGGC<br>ATTCACCGACGTTAAGGTCCCCCCAACTACAACTCCCTCTCCCCCGCCGGTGAGCAAAGGTGAGGCCGTGATCAAGGAATTCATGCGTTTCAAGGTCC<br>ACATGGAAGGATCAATGAACGGACACGAATTCGAGATAGAAGGGGAAGGCGAAGGTCGTCCATACGAAGGTACGCAAACAGCAAAACTGAAGGTGACG |
| MUC2_R3 | SPPPTSTTTLP | CCGATGGCCGCAAACTACCTTAAGAACCAGCCCATGTACGTGTTTCGGAAAACTGAGCTTAAACACAGCAAGACAGAGTTAAATTTCAAGGAGTGGCAAAAGGC<br>ATTCACCGACGTTAAGTCACCACCGCCTACATCAACAACAACTCTGCCAGTGAGCAAAGGTGAGGCCGTGATCAAGGAATTCATGCGTTTCAAGGTCC<br>ACATGGAAGGATCAATGAACGGACACGAATTCGAGATAGAAGGGGAAGGCGAAGGTCGTCCATACGAAGGTACGCAAACAGCAAAACTGAAGGTGACG |
| MUC5AC_R1 | PPTTSTTSAPP | CCGATGGCCGCAAACTACCTTAAGAACCAGCCCATGTACGTGTTTCGGAAAACTGAGCTTAAACACAGCAAGACAGAGTTAAATTTCAAGGAGTGGCAAAAGGC<br>ATTCACCGACGTTAAGCCGCCGACGACATCGACGACATCTGCTCCCCCCGTGAGCAAAGGTGAGGCCGTGATCAAGGAATTCATGCGTTTCAAGGTCC<br>ACATGGAAGGATCAATGAACGGACACGAATTCGAGATAGAAGGGGAAGGCGAAGGTCGTCCATACGAAGGTACGCAAACAGCAAAACTGAAGGTGACG |
| MUC5AC_R2 | PVPATTVGPVP | CCGATGGCCGCAAACTACCTTAAGAACCAGCCCATGTACGTGTTTCGGAAAACTGAGCTTAAACACAGCAAGACAGAGTTAAATTTCAAGGAGTGGCAAAAGGC<br>ATTCACCGACGTTAAGCCCGTTCCCGCCACAACTGTTGGGCCTGTTCCTGTGAGCAAAGGTGAGGCCGTGATCAAGGAATTCATGCGTTTCAAGGTCC<br>ACATGGAAGGATCAATGAACGGACACGAATTCGAGATAGAAGGGGAAGGCGAAGGTCGTCCATACGAAGGTACGCAAACAGCAAAACTGAAGGTGACG |
| MUC5B_R1 | GSTATPSSTPG | CCGATGGCCGCAAACTACCTTAAGAACCAGCCCATGTACGTGTTTCGGAAAACTGAGCTTAAACACAGCAAGACAGAGTTAAATTTCAAGGAGTGGCAAAAGGC<br>ATTCACCGACGTTAAGGGATCGACCGCCACACCGTCTTCAACACCTGGGGTGAGCAAAGGTGAGGCCGTGATCAAGGAATTCATGCGTTTCAAGGTCC<br>ACATGGAAGGATCAATGAACGGACACGAATTCGAGATAGAAGGGGAAGGCGAAGGTCGTCCATACGAAGGTACGCAAACAGCAAAACTGAAGGTGACG |
| MUC5B_R2 | VLTTTATTPTP | CCGATGGCCGCAAACTACCTTAAGAACCAGCCCATGTACGTGTTTCGGAAAACTGAGCTTAAACACAGCAAGACAGAGTTAAATTTCAAGGAGTGGCAAAAGGC<br>ATTCACCGACGTTAAGGTACTTACTACTACGGCGACGACACCAACCCCTGTGAGCAAAGGTGAGGCCGTGATCAAGGAATTCATGCGTTTCAAGGTCC<br>ACATGGAAGGATCAATGAACGGACACGAATTCGAGATAGAAGGGGAAGGCGAAGGTCGTCCATACGAAGGTACGCAAACAGCAAAACTGAAGGTGACG |
| IgA1_h | SPSTPPTPSPS | CCGATGGCCGCAAACTACCTTAAGAACCAGCCCATGTACGTGTTTCGGAAAACTGAGCTTAAACACAGCAAGACAGAGTTAAATTTCAAGGAGTGGCAAAAGGC<br>ATTCACCGACGTTAAGTCACCGAGTACACCTCCTACACCGAGCCCTTCGGTGAGCAAAGGTGAGGCCGTGATCAAGGAATTCATGCGTTTCAAGGTCC<br>ACATGGAAGGATCAATGAACGGACACGAATTCGAGATAGAAGGGGAAGGCGAAGGTCGTCCATACGAAGGTACGCAAACAGCAAAACTGAAGGTGACG |
| MUC4 | STGHATPLPVT | CCGATGGCCGCAAACTACCTTAAGAACCAGCCCATGTACGTGTTTCGGAAAACTGAGCTTAAACACAGCAAGACAGAGTTAAATTTCAAGGAGTGGCAAAAGGC<br>ATTCACCGACGTTAAGTCGACAGGCCATGCAACACCCCTTCCAGTAACAGTGAGCAAAGGTGAGGCCGTGATCAAGGAATTCATGCGTTTCAAGGTCC<br>ACATGGAAGGATCAATGAACGGACACGAATTCGAGATAGAAGGGGAAGGCGAAGGTCGTCCATACGAAGGTACGCAAACAGCAAAACTGAAGGTGACG |
| MUC3A/B | TSSITTTETTS | CCGATGGCCGCAAACTACCTTAAGAACCAGCCCATGTACGTGTTTCGGAAAACTGAGCTTAAACACAGCAAGACAGAGTTAAATTTCAAGGAGTGGCAAAAGGC<br>ATTCACCGACGTTAAGACAAGCTCGATTACCACAACTGAGACGACATCGGTGAGCAAAGGTGAGGCCGTGATCAAGGAATTCATGCGTTTCAAGGTCC<br>ACATGGAAGGATCAATGAACGGACACGAATTCGAGATAGAAGGGGAAGGCGAAGGTCGTCCATACGAAGGTACGCAAACAGCAAAACTGAAGGTGACG |
| MUC7_R1 | TTAAPPTPSAT | CCGATGGCCGCAAACTACCTTAAGAACCAGCCCATGTACGTGTTTCGGAAAACTGAGCTTAAACACAGCAAGACAGAGTTAAATTTCAAGGAGTGGCAAAAGGC<br>ATTCACCGACGTTAAGACGACCGCTGCCCCCCCGACACCGTCGGCAACAGTGAGCAAAGGTGAGGCCGTGATCAAGGAATTCATGCGTTTCAAGGTCC<br>ACATGGAAGGATCAATGAACGGACACGAATTCGAGATAGAAGGGGAAGGCGAAGGTCGTCCATACGAAGGTACGCAAACAGCAAAACTGAAGGTGACG |
| MUC7_R2 | PSATTPAPPSS | CCGATGGCCGCAAACTACCTTAAGAACCAGCCCATGTACGTGTTTCGGAAAACTGAGCTTAAACACAGCAAGACAGAGTTAAATTTCAAGGAGTGGCAAAAGGC<br>ATTCACCGACGTTAAGCCTTCAGCAACTACACCGGCCCCACCTTCGTCTGTGAGCAAAGGTGAGGCCGTGATCAAGGAATTCATGCGTTTCAAGGTCC<br>ACATGGAAGGATCAATGAACGGACACGAATTCGAGATAGAAGGGGAAGGCGAAGGTCGTCCATACGAAGGTACGCAAACAGCAAAACTGAAGGTGACG |
| MUC8_S1 | PLQEGTPGSRV | CCGATGGCCGCAAACTACCTTAAGAACCAGCCCATGTACGTGTTTCGGAAAACTGAGCTTAAACACAGCAAGACAGAGTTAAATTTCAAGGAGTGGCAAAAGGC<br>ATTCACCGACGTTAAGCCATTGCAAGAAGGCACACCTGGATCTCGTGTGGTGAGCAAAGGTGAGGCCGTGATCAAGGAATTCATGCGTTTCAAGGTCC<br>ACATGGAAGGATCAATGAACGGACACGAATTCGAGATAGAAGGGGAAGGCGAAGGTCGTCCATACGAAGGTACGCAAACAGCAAAACTGAAGGTGACG |
| MUC8_S2 | VHELPTSSPGG | CCGATGGCCGCAAACTACCTTAAGAACCAGCCCATGTACGTGTTTCGGAAAACTGAGCTTAAACACAGCAAGACAGAGTTAAATTTCAAGGAGTGGCAAAAGGC<br>ATTCACCGACGTTAAGGTGCACGAGTTGCCGACCAGCTCCCCTGGTGGCGTGAGCAAAGGTGAGGCCGTGATCAAGGAATTCATGCGTTTCAAGGTCC<br>ACATGGAAGGATCAATGAACGGACACGAATTCGAGATAGAAGGGGAAGGCGAAGGTCGTCCATACGAAGGTACGCAAACAGCAAAACTGAAGGTGACG |
| MUC10 | TTDSTTPAPTT | CCGATGGCCGCAAACTACCTTAAGAACCAGCCCATGTACGTGTTTCGGAAAACTGAGCTTAAACACAGCAAGACAGAGTTAAATTTCAAGGAGTGGCAAAAGGC<br>ATTCACCGACGTTAAGACTACTGATTCCACCACGCCTGCGCCAACTACAGTGAGCAAAGGTGAGGCCGTGATCAAGGAATTCATGCGTTTCAAGGTCC<br>ACATGGAAGGATCAATGAACGGACACGAATTCGAGATAGAAGGGGAAGGCGAAGGTCGTCCATACGAAGGTACGCAAACAGCAAAACTGAAGGTGACG |
| MUC11/12 | SSPGSTHTTLS | CCGATGGCCGCAAACTACCTTAAGAACCAGCCCATGTACGTGTTTCGGAAAACTGAGCTTAAACACAGCAAGACAGAGTTAAATTTCAAGGAGTGGCAAAAGGC<br>ATTCACCGACGTTAAGTCCTCCCCCGGATCTACTCACACTACACTGTCAGTGAGCAAAGGTGAGGCCGTGATCAAGGAATTCATGCGTTTCAAGGTCC<br>ACATGGAAGGATCAATGAACGGACACGAATTCGAGATAGAAGGGGAAGGCGAAGGTCGTCCATACGAAGGTACGCAAACAGCAAAACTGAAGGTGACG |
| MUC16 | SVPTTSTPGTS | CCGATGGCCGCAAACTACCTTAAGAACCAGCCCATGTACGTGTTTCGGAAAACTGAGCTTAAACACAGCAAGACAGAGTTAAATTTCAAGGAGTGGCAAAAGGC<br>ATTCACCGACGTTAAGAGTGTGCCGACAACCTCGACACCAGGGACAAGCGTGAGCAAAGGTGAGGCCGTGATCAAGGAATTCATGCGTTTCAAGGTCC<br>ACATGGAAGGATCAATGAACGGACACGAATTCGAGATAGAAGGGGAAGGCGAAGGTCGTCCATACGAAGGTACGCAAACAGCAAAACTGAAGGTGACG |
| MUC19 | STTVAPGSTTV | CCGATGGCCGCAAACTACCTTAAGAACCAGCCCATGTACGTGTTTCGGAAAACTGAGCTTAAACACAGCAAGACAGAGTTAAATTTCAAGGAGTGGCAAAAGGC<br>ATTCACCGACGTTAAGTCGACTACTGTTGCTCCAGGTAGCACAACGGTAGTGAGCAAAGGTGAGGCCGTGATCAAGGAATTCATGCGTTTCAAGGTCC<br>ACATGGAAGGATCAATGAACGGACACGAATTCGAGATAGAAGGGGAAGGCGAAGGTCGTCCATACGAAGGTACGCAAACAGCAAAACTGAAGGTGACG |
| ASF | AEAPTAVPDKG | CCGATGGCCGCAAACTACCTTAAGAACCAGCCCATGTACGTGTTTCGGAAAACTGAGCTTAAACACAGCAAGACAGAGTTAAATTTCAAGGAGTGGCAAAAGGC<br>ATTCACCGACGTTAAGGCAGAGGCCCCGACTGCCGTGCCTGATAAAGGCGTGAGCAAAGGTGAGGCCGTGATCAAGGAATTCATGCGTTTCAAGGTCC<br>ACATGGAAGGATCAATGAACGGACACGAATTCGAGATAGAAGGGGAAGGCGAAGGTCGTCCATACGAAGGTACGCAAACAGCAAAACTGAAGGTGACG |
| Glycoprotein Iba | PAPSPTTPEPT | CCGATGGCCGCAAACTACCTTAAGAACCAGCCCATGTACGTGTTTCGGAAAACTGAGCTTAAACACAGCAAGACAGAGTTAAATTTCAAGGAGTGGCAAAAGGC<br>ATTCACCGACGTTAAGCCGGCACCCTCACCCACTACTCCGGAGCCAACTGTGAGCAAAGGTGAGGCCGTGATCAAGGAATTCATGCGTTTCAAGGTCC<br>ACATGGAAGGATCAATGAACGGACACGAATTCGAGATAGAAGGGGAAGGCGAAGGTCGTCCATACGAAGGTACGCAAACAGCAAAACTGAAGGTGACG |
| MUC6_R1 | TTTYPTPSHPQ | CCGATGGCCGCAAACTACCTTAAGAACCAGCCCATGTACGTGTTTCGGAAAACTGAGCTTAAACACAGCAAGACAGAGTTAAATTTCAAGGAGTGGCAAAAGGC<br>ATTCACCGACGTTAAGACGACGACGTATCCCACCCCGAGTCACCCTCAAGTGAGCAAAGGTGAGGCCGTGATCAAGGAATTCATGCGTTTCAAGGTCC<br>ACATGGAAGGATCAATGAACGGACACGAATTCGAGATAGAAGGGGAAGGCGAAGGTCGTCCATACGAAGGTACGCAAACAGCAAAACTGAAGGTGACG |
| MUC17_R1 | SPTNSSPTTAE | CCGATGGCCGCAAACTACCTTAAGAACCAGCCCATGTACGTGTTTCGGAAAACTGAGCTTAAACACAGCAAGACAGAGTTAAATTTCAAGGAGTGGCAAAAGGC<br>ATTCACCGACGTTAAGTCACCAACCAATTCGTCTCCGACCACTGCTGAGGTGAGCAAAGGTGAGGCCGTGATCAAGGAATTCATGCGTTTCAAGGTCC<br>ACATGGAAGGATCAATGAACGGACACGAATTCGAGATAGAAGGGGAAGGCGAAGGTCGTCCATACGAAGGTACGCAAACAGCAAAACTGAAGGTGACG |
| Primer 1 | | TTTGCGGCCATCGGCTTAGC |
| Primer 2 | | AAACTGAAGGTGACGAAAGGTGGCC |

**Supplementary Table 2.** Properties of *Bacteroides caccae* M60 peptidases.

| *B. caccae* M60 | Locus Tag (new assembly) | Locus Tag (original assembly) | Molecular Weight (g/mol) | Extinction Coefficient ($M^{-1}$ $cm^{-1}$) | *E. coli* expression strain |
|---|---|---|---|---|---|
| BcM60A | CGC64_02005 | BACCAC_00161 | 61278.36 | 107635 | BL21 (DE3) |
| BcM60B | CGC64_02920 | BACCAC_01246 | 53885.47 | 79565 | BL21 (DE3) |
| BcM60C | CGC64_03500 | BACCAC_01368 | 59409.12 | 112565 | BL21 (DE3) |
| BcM60D | CGC64_03540 | BACCAC_01376 | 61326.58 | 92750 | Shuffle T7 |
| BcM60E | CGC64_03545 | BACCAC_01377 | 61439.74 | 117730 | BL21 (DE3) |
| BcM60F | CGC64_04090 | BACCAC_01495 | 59371.78 | 87920 | BL21 (DE3) |
| BcM60G | CGC64_04095 | BACCAC_01496 | 49739.70 | 84355 | BL21 (DE3) |
| BcM60H | CGC64_07210 | BACCAC_01782 | 62521.22 | 127130 | BL21 (DE3) |
| BcM60I | CGC64_07485 | BACCAC_01840 | 64460.95 | 118065 | Shuffle T7 |
| BcM60J | CGC64_07490 | BACCAC_01841 | 65174.52 | 138910 | Shuffle T7 |
| BcM60K | CGC64_07580 | BACCAC_01859 | 60926.43 | 97555 | BL21 (DE3) |
| BcM60L | CGC64_08375 | BACCAC_03612 | 60932.89 | 107065 | BL21 (DE3) |
| BcM60M | CGC64_07535 | BACCAC_01850 | | | BL21 (DE3) |
| BcM60N | CGC64_14820 | BACCAC_02574 | N/A | N/A | N/A |
| BcM60O | CGC64_14820 | BACCAC_02574 | N/A | N/A | N/A |
| BcM60P | CGC64_16100 | BACCAC_02897 | N/A | N/A | N/A |

**Supplementary Table 3.** Crystallization conditions of *Bacteroides caccae* M60 peptidases.

| Protein | Notes | [Protein] | Crystallization condition | Cryoprotectant |
|---|---|---|---|---|
| BcM60B | His tag removed | 12.5 | 4% Tascimate pH4.0<br>15% PEG3350<br>Cryo 20% EG | 20% ethylene glycol |
| BcM60C<br>WT and E665Q mutant | His tag removed | 29 | 0.2M $(NH_4)_2$ $SO_4$<br>0.1M Na citrate pH5.6<br>21% PEG4000 | 20% ethylene glycol |
| BcM60F | His tag removed | 10 | 0.1M Na citrate tribasic dihydrate pH5.6<br>20% 2-Propanol<br>20% PEG4K | 20% ethylene glycol |
| BcM60K<br>WT and E664A mutant | His tag removed | 15.3 | 0.1M Bis-Tris pH6.0<br>20% PEG3350 | 20% ethylene glycol |

**Supplementary Table 4. X-ray** Data collection and structure statistics. Values for highest resolution shells are shown in parenthesis.

| | BcM60K | BcM60K C2 | BcM60K 6SC1 |
|---|---|---|---|
| ***Data Collection*** | | | |
| Beamline | Home Beam | Home Beam | Home Beam |
| Wavelength (Å) | 1.54178 | 1.54178 | 1.54178 |
| Space Group | $P2_1$ | $P2_1$ | $P2_1$ |
| Cell Dimensions | | | |
| *a, b, c* (Å) | 46.9, 64.9, 93.4 | 46.9, 64.9, 93.6 | 46.5, 65.1, 93.2 |
| $\alpha, \beta, \gamma$ (°) | 90.0, 100.8, 90.0 | 90.0, 100.8, 90.0 | 90.0, 100.6, 90.0 |
| Resolution (Å) | 30.00-2.10 (2.14-2.10) | 30.00-2.50 (2.54-2.50) | 30.00-2.50 (2.54-2.50) |
| $R_{meas}$ | 0.092 (0.288) | 0.144 (0.709) | 0.078 (0.289) |
| $R_{pim}$ | 0.047 (0.199) | 0.071 (0.410) | 0.042 (0.183) |
| CC1/2 | 0.983 (0.894) | 0.987 (0.631) | 0.994 (0.912) |
| $\langle I/\sigma I \rangle$ | 11.6 (2.5) | 8.9 (1.7) | 14.4 (3.3) |
| Completeness (%) | 98.8 (94.2) | 99.5 (95.2) | 99.1 (92.8) |
| Redundancy | 2.9 (2.3) | 3.8 (2.6) | 2.9 (2.0) |
| No. of reflections | 92946 | 73443 | 53428 |
| No. Unique | 31868 (1503) | 19286 (885) | 19338 (883) |
| ***Refinement*** | | | |
| Resolution (Å) | 2.10 | 2.5 | 2.5 |
| $R_{work}/R_{free}$ | 0.22/0.25 | 0.02/0.24 | 0.22/0.28 |
| No. of atoms | | | |
| Protein | 4221 | 4187 | 4193 |
| Ligand | 1 (ZN), 28 (EDO), 14 (BTB) | 1 (ZN), 47 (C2) | 1 (ZN), 20 (EDO), 52 (6SC1) |
| Water | 263 | 200 | 92 |
| *B*-factors | | | |
| Protein | 29.3 | 41.5 | 40.0 |
| Ligand | 29.9 (ZN), 30.0 (EDO), 46.9 (BTB) | 44.1 (ZN), 50.1 (C2) | 40.2 (ZN), 38.9 (EDO), 45.8 (6SC1) |
| Water | 27.9 | 37.9 | 33.5 |
| r.m.s.d | | | |
| Bond lengths (Å) | 0.001 | 0.003 | 0.002 |
| Bond angles (°) | 0.416 | 0.539 | 0.469 |
| Ramachandran (%) | | | |
| Preferred | 94.8 | 95.6 | 94.5 |
| Allowed | 5.2 | 4.4 | 5.5 |
| Disallowed | 0.0 | 0.0 | 0.0 |

| | BcM60K_E664A Muc5AC peptide | BcM60F | BcM60B |
|---|---|---|---|
| ***Data Collection*** | | | |
| Beamline | Home Beam | Home Beam | Home Beam |
| Wavelength (Å) | 1.54178 | 1.54178 | 1.54178 |
| Space Group | $P2_1$ | P21 | P21 |
| Cell Dimensions | | | |
| *a, b, c* (Å) | 46.9, 65.0, 93.6 | 78.3, 96.7, 85.9 | 48.1, 109.4, 96.8 |
| $\alpha, \beta, \gamma$ (°) | 90.0, 100.9, 90.0 | 90.0, 114.6, 90.0 | 90.0, 94.8, 90.0 |
| Resolution (Å) | 25.00-2.20 (2.24-2.20) | 30.00-1.80 (1.83-1.80) | 30.00-1.80 (1.83-1.80) |
| $R_{meas}$ | 0.106 (0.525) | 0.055 (0.166) | 0.070 (0.251) |
| $R_{pim}$ | 0.052 (0.290) | 0.025 (0.100) | 0.034 (0.162) |
| CC1/2 | 0.919 (0.997) | 0.998 (0.959) | 0.997 (0.906) |
| $\langle I/\sigma I \rangle$ | 13.1 (2.4) | 27.4 (6.2) | 19.4 (4.2) |
| Completeness (%) | 99.6 (96.2) | 99.0 (97.8) | 100.0 (99.6) |
| Redundancy | 3.8 (2.5) | 3.9 (2.9) | 4.0 (2.9) |
| No. of reflections | 106679 | 413212 | 371777 |
| No. Unique | 28358 (1401) | 106768 (5256) | 92252 (4571) |
| ***Refinement*** | | | |
| Resolution (Å) | 2.20 | 1.80 | 1.80 |
| $R_{work}/R_{free}$ | 0.18/0.22 | 0.15/0.18 | 0.17/0.20 |
| No. of atoms | | | |
| Protein | 4254 | 4122 (A), 4104 (B) | 3771 (A), 3829 (B) |
| Ligand | 1 (ZN), 47 (peptide) | 48 (EDO) | 2 (ZN), 40 EDO), 21 (MLA), 1 (CA), 15 (FMT), 16 (SIN) |
| Water | 407 | 1085 | 714 |
| *B*-factors | | | |
| Protein | | 18.7 (A), 17.7 (B) | 20.5 (A), 18.0 (B) |
| Ligand | 35.4 (ZN), 59.6 (peptide) | 29.2 (EDO) | 28.5 (ZN), 25.8 (EDO),31.0 (MLA), 22.8 (CA), 27.6 (FMT), 33.9 (SIN) |
| Water | 38.4 | 27.0 | 26.4 |
| r.m.s.d | | | |
| Bond lengths (Å) | 0.008 | 0.007 | 0.011 |
| Bond angles (°) | 0.870 | 0.902 | 1.756 |
| Ramachandran (%) | | | |
| Preferred | 95.0 | 97.7 | 97.9 |
| Allowed | 5.0 | 2.3 | 2.1 |
| Disallowed | 0.0 | 0.0 | 0.0 |

| | BcM60C | BcM60C 6SC1 | BcM60C_E665Q C2 |
|---|---|---|---|
| ***Data Collection*** | | | |
| Beamline | Home Beam | Home Beam | Home Beam |
| Wavelength (Å) | 1.54178 | 1.54178 | 1.54178 |
| Space Group | $P4_12_12$ | $P4_12_12$ | $P4_12_12$ |
| Cell Dimensions | | | |
| *a, b, c* (Å) | 118.6, 118.6, 90.9 | 118.5, 118.5, 90.8 | 118.4, 118.4, 88.2 |
| $\alpha$, $\beta$, $\gamma$ (º) | 90.0, 90.0, 90.0 | 90.0, 90.0, 90.0 | 90.0, 90.0, 90.0 |
| Resolution (Å) | 30.00-1.55 (1.58-1.55) | 30.00-1.55 (1.58-1.55) | 25.00-2.15 (2.19-2.15) |
| $R_{merge}$ | 0.089 (0.569) | 0.056 (0.384) | 0.087 (0.422) |
| $R_{pim}$ | 0.026 (0.263) | 0.022 (0.267) | 0.039 (0.307) |
| CC1/2 | 0.997 (0.855) | 0.997 (0.856) | 0.998 (0.778) |
| $\langle I/\sigma I \rangle$ | 27.5 (2.4) | 27.8 (2.3) | 16.1 (2.1) |
| Completeness (%) | 99.8 (99.9) | 99.8 (99.7) | 98.5 (87.1) |
| Redundancy | 7.8 (5.0) | 4.2 (2.7) | 4.1 (2.1) |
| No. of reflections | 730718 | 396537 | 140286 |
| No. Unique | 93992 (4620) | 93659 (4620) | 34134 (1451) |
| ***Refinement*** | | | |
| Resolution (Å) | 1.55 | 1.55 | 2.15 |
| $R_{work}/R_{free}$ | 0.18/0.21 | 0.18/0.21 | 0.21/0.26 |
| No. of atoms | | | |
| Protein | 4173 | 4182 | 4038 |
| Ligand | 1 (ZN), 15 (SO4), 16 (EDO) | 1 (ZN), 15 (SO4), 16 (EDO), 52 (STAg) | 1 (ZN), 10 (SO4), 24 (EDO), 47 (Thr_Core2) |
| Water | 678 | 674 | 230 |
| *B*-factors | | | |
| Protein | 17.2 | 17.1 | 27.5 |
| Ligand | 16.6 (ZN), 37.9 (SO4), 28.4 (EDO), | 15.0 (ZN), 28.2 (SO4), 28.7 (EDO), 32.3 (STAg) | 24.4 (ZN), 45.2 (SO4), 39.6 (SO4), 35.8 (Thr_Core2) |
| Water | 28.0 | 27.9 | 27.6 |
| r.m.s.d | | | |
| Bond lengths (Å) | 0.009 | 0.010 | 0.002 |
| Bond angles (°) | 1.051 | 1.081 | 0.453 |
| Ramachandran (%) | | | |
| Preferred | 96.9 | 97.1 | 95.3 |
| Allowed | 3.1 | 2.9 | 4.7 |
| Disallowed | 0.0 | 0.0 | 0.0 |

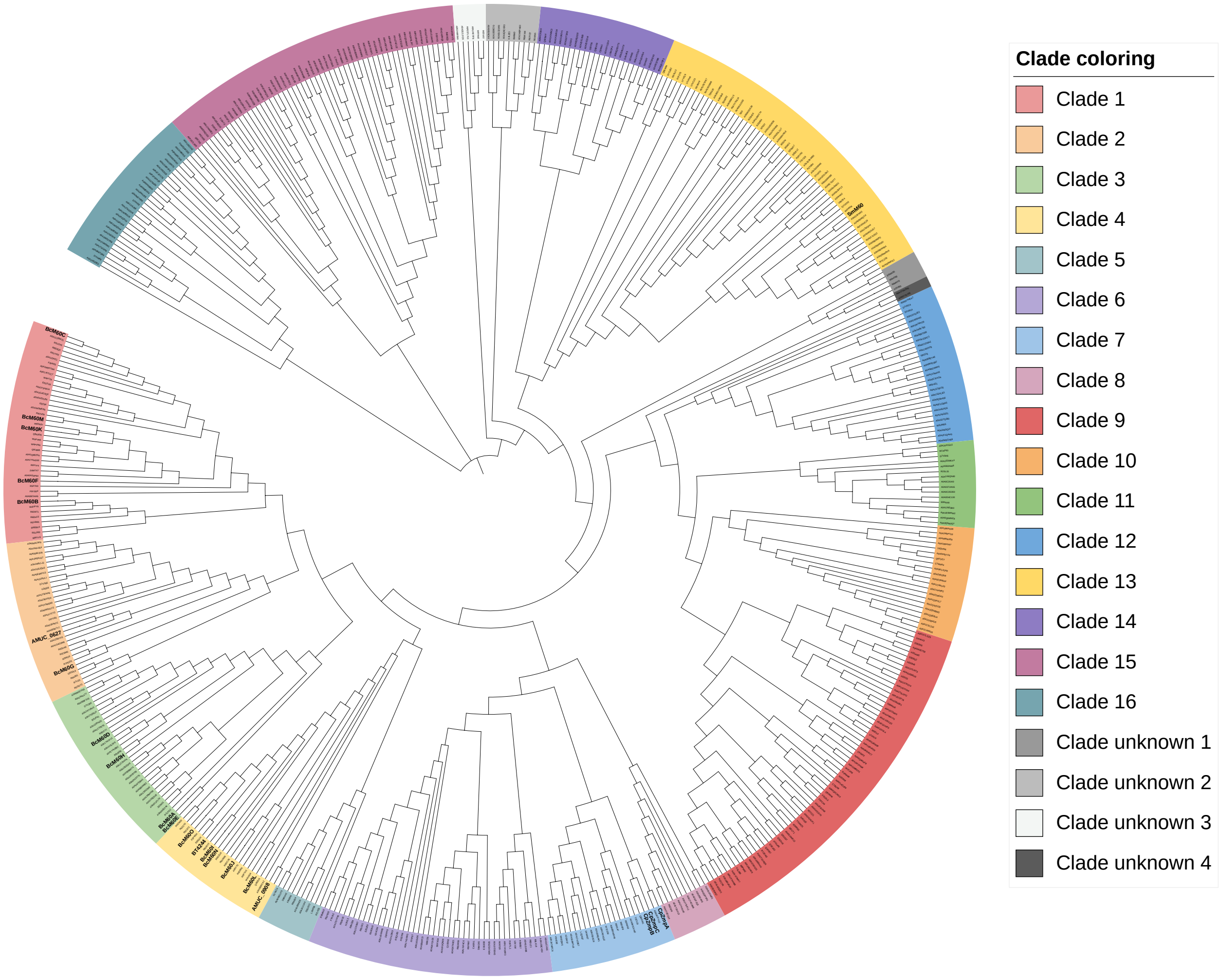

**Supplementary Figure 1.** Complete phylogenetic tree of peptidase_M60 proteins from Pfam entry 13402 (Pfam version 32). The *B. caccae* proteins are labeled black, BcM60C and McM60K, which are the primary topics of this study are in green. Entries with prior characterization are labeled in blue.

A

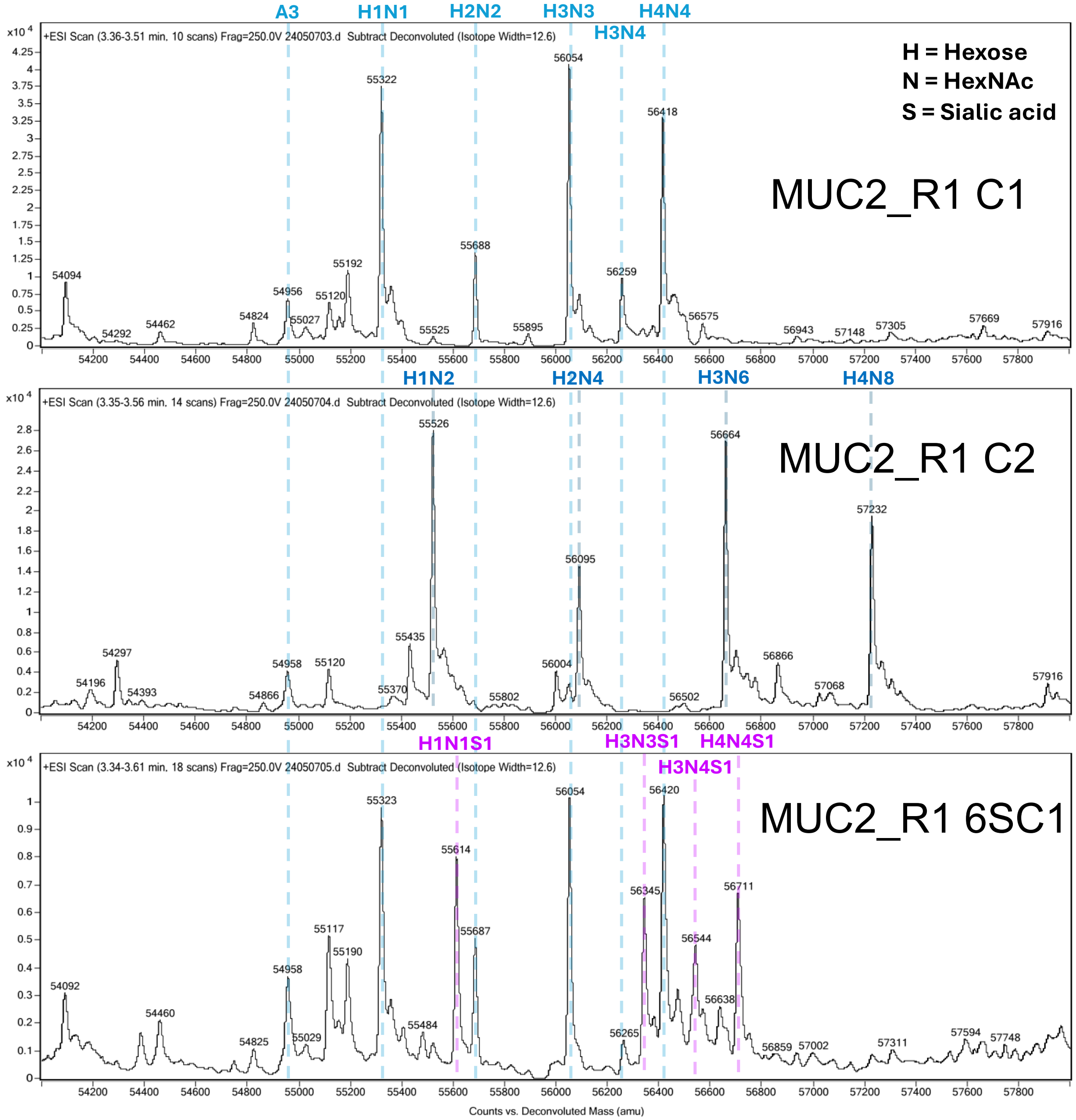
A3
H1N1
H2N2
H3N3
H4N4
H3N4
H = Hexose
N = HexNAc
S = Sialic acid
MUC2_R1 C1
+ESI Scan (3.36-3.51 min. 10 scans) Frag=250.0V 24050703.d Subtract Deconvoluted (Isotope Width=12.6)
H1N2
H2N4
H3N6
H4N8
MUC2_R1 C2
+ESI Scan (3.35-3.56 min. 14 scans) Frag=250.0V 24050704.d Subtract Deconvoluted (Isotope Width=12.6)
H1N1S1
H3N3S1
H4N4S1
H3N4S1
MUC2_R1 6SC1
+ESI Scan (3.34-3.61 min. 18 scans) Frag=250.0V 24050705.d Subtract Deconvoluted (Isotope Width=12.6)
Counts vs. Deconvoluted Mass (amu)

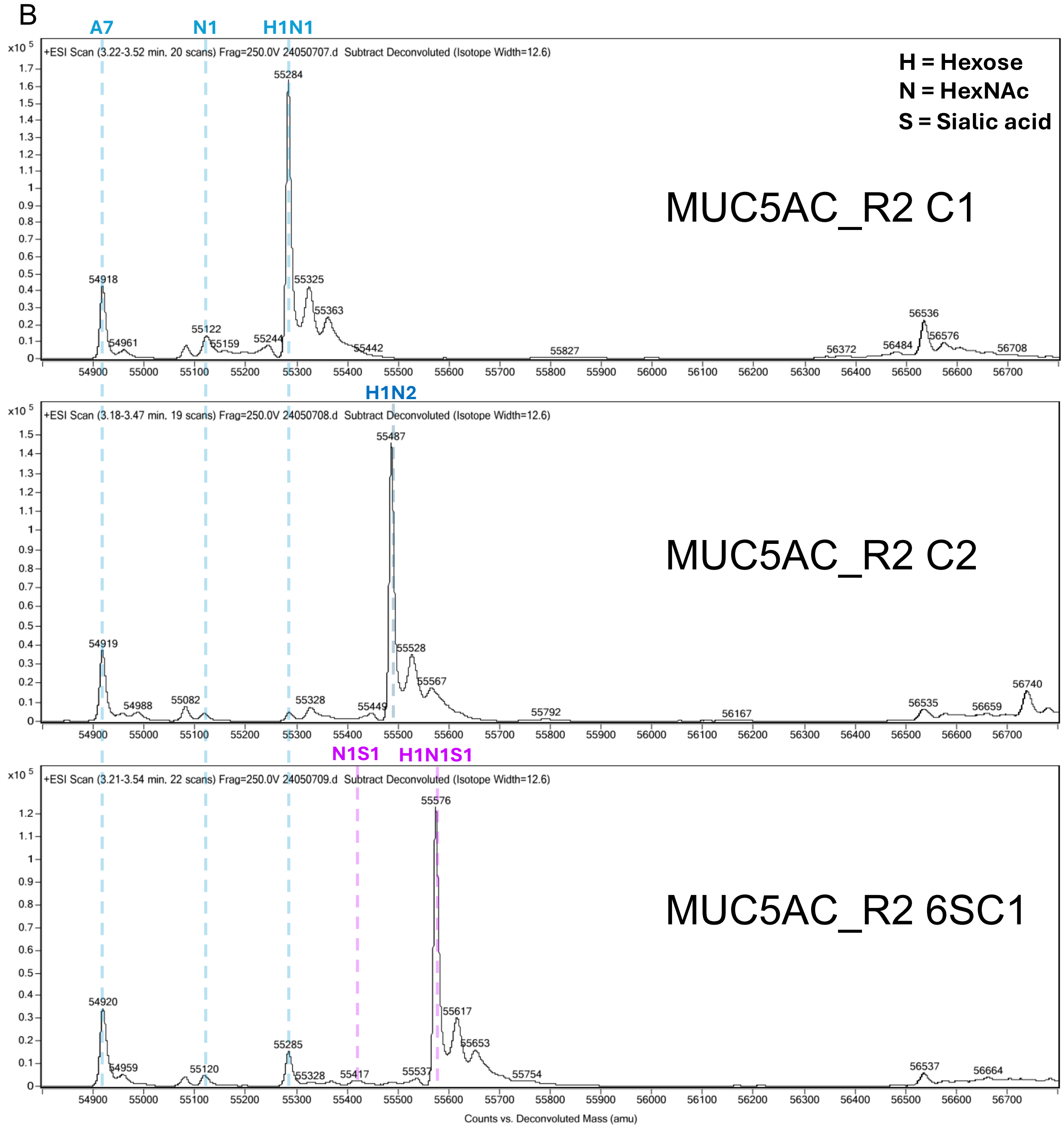
B
A7
N1
H1N1
H1N2
N1S1
H1N1S1
H = Hexose
N = HexNAc
S = Sialic acid
MUC5AC_R2 C1
MUC5AC_R2 C2
MUC5AC_R2 6SC1
+ESI Scan (3.22-3.52 min, 20 scans) Frag=250.0V 24050707.d Subtract Deconvoluted (Isotope Width=12.6)
+ESI Scan (3.18-3.47 min, 19 scans) Frag=250.0V 24050708.d Subtract Deconvoluted (Isotope Width=12.6)
+ESI Scan (3.21-3.54 min, 22 scans) Frag=250.0V 24050709.d Subtract Deconvoluted (Isotope Width=12.6)
Counts vs. Deconvoluted Mass (amu)

C

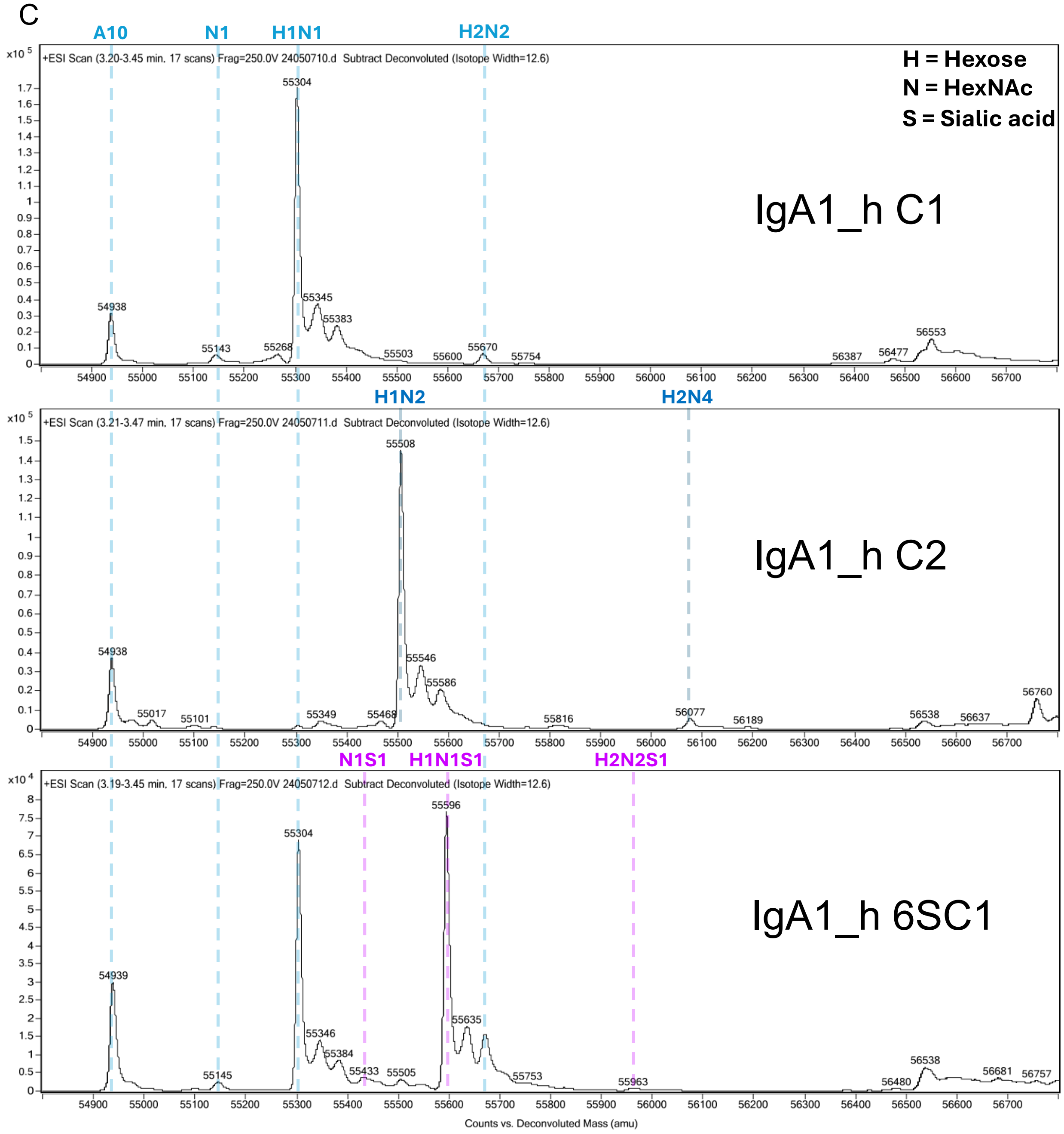
A10
N1
H1N1
H2N2
H1N2
H2N4
N1S1
H1N1S1
H2N2S1
H = Hexose
N = HexNAc
S = Sialic acid
IgA1_h C1
IgA1_h C2
IgA1_h 6SC1
+ESI Scan (3.20-3.45 min, 17 scans) Frag=250.0V 24050710.d Subtract Deconvoluted (Isotope Width=12.6)
+ESI Scan (3.21-3.47 min, 17 scans) Frag=250.0V 24050711.d Subtract Deconvoluted (Isotope Width=12.6)
+ESI Scan (3.19-3.45 min, 17 scans) Frag=250.0V 24050712.d Subtract Deconvoluted (Isotope Width=12.6)
x10 5
x10 4
54938
55143
55268
55304
55345
55383
55503
55600
55670
55754
56387
56477
56553
55017
55101
55349
55468
55508
55546
55586
55816
56077
56189
56538
56637
56760
54939
55145
55346
55384
55433
55505
55596
55635
55753
55963
56480
56681
56757
Counts vs. Deconvoluted Mass (amu)

D

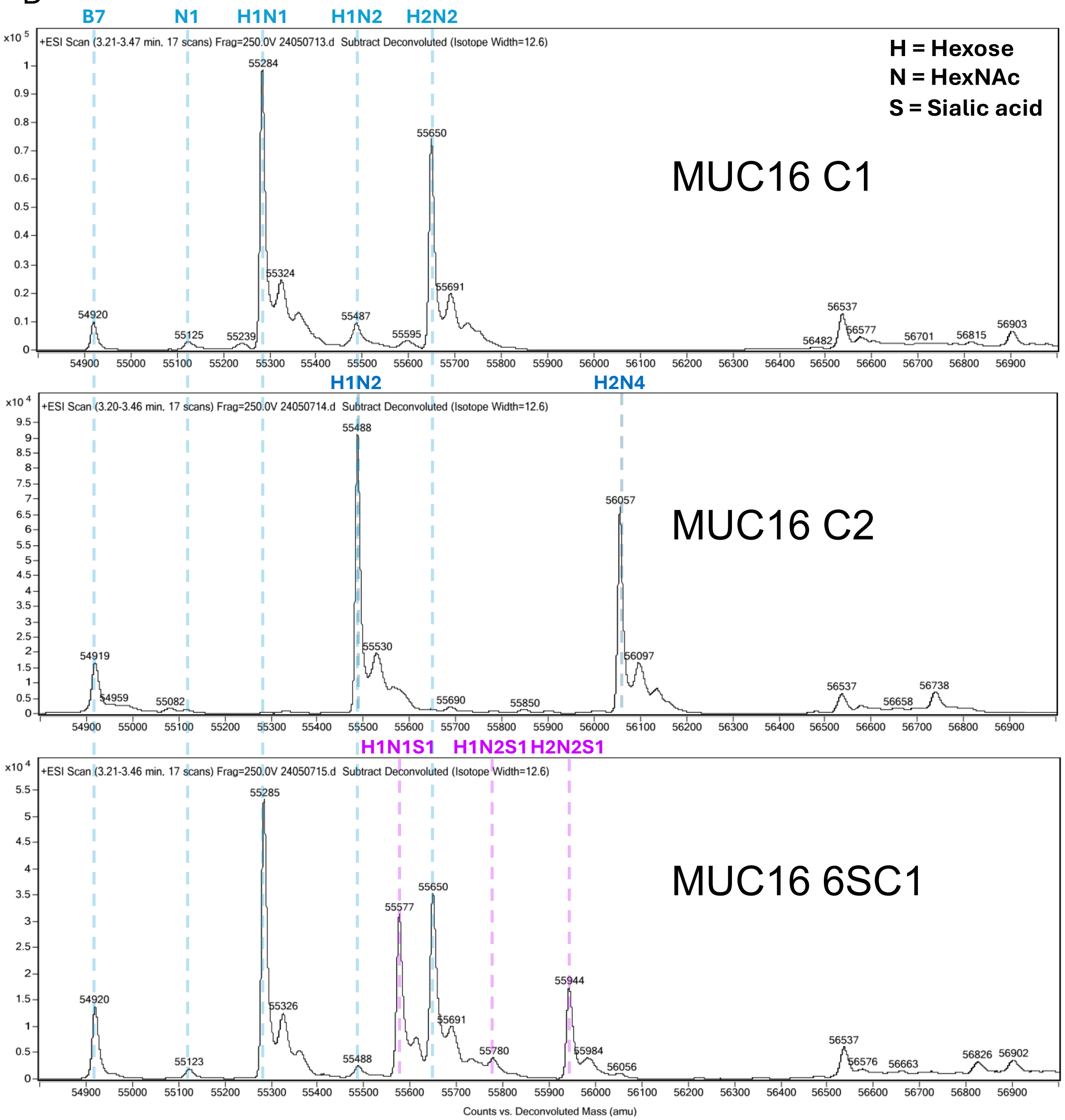

**Supplementary Figure 2.** Intact mass spectrometry analysis of four FRET substrates modified with C1, C2, and 6SC1. A) MUC2_R1 substrate. B) MUC5AC_R2 substrate. C) IgA1_h substrate. D) MUC16 substrate. Linker identities are provided in each panel.

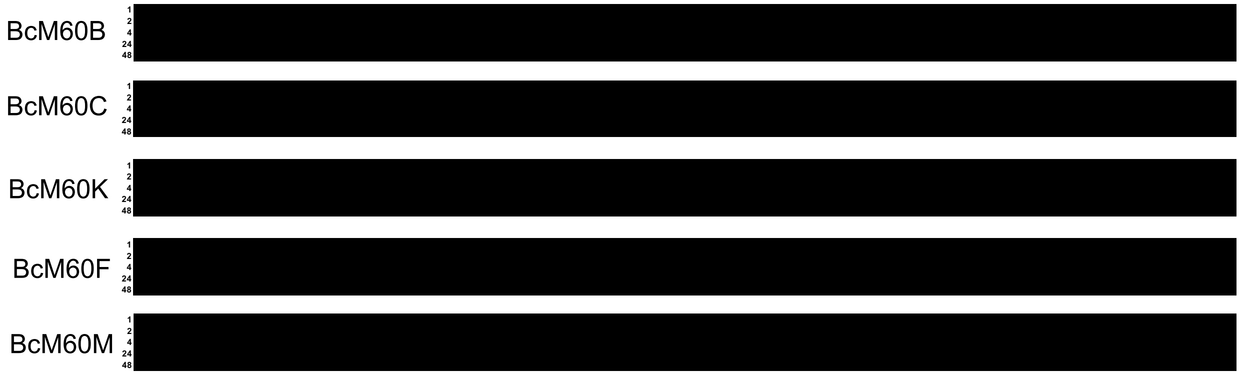

**Supplementary Figure 3.** FRET screens using unglycosylated substrates. Heat maps show FRET values calculated by the ratio-of-ratios approach. The coloring is ramped from a low threshold of 0.2 to a high of 1.0. Numbers just to the left of the heat maps indicate the time of incubation in hours. The observation that all of the samples are black indicates a lack of significant activity.

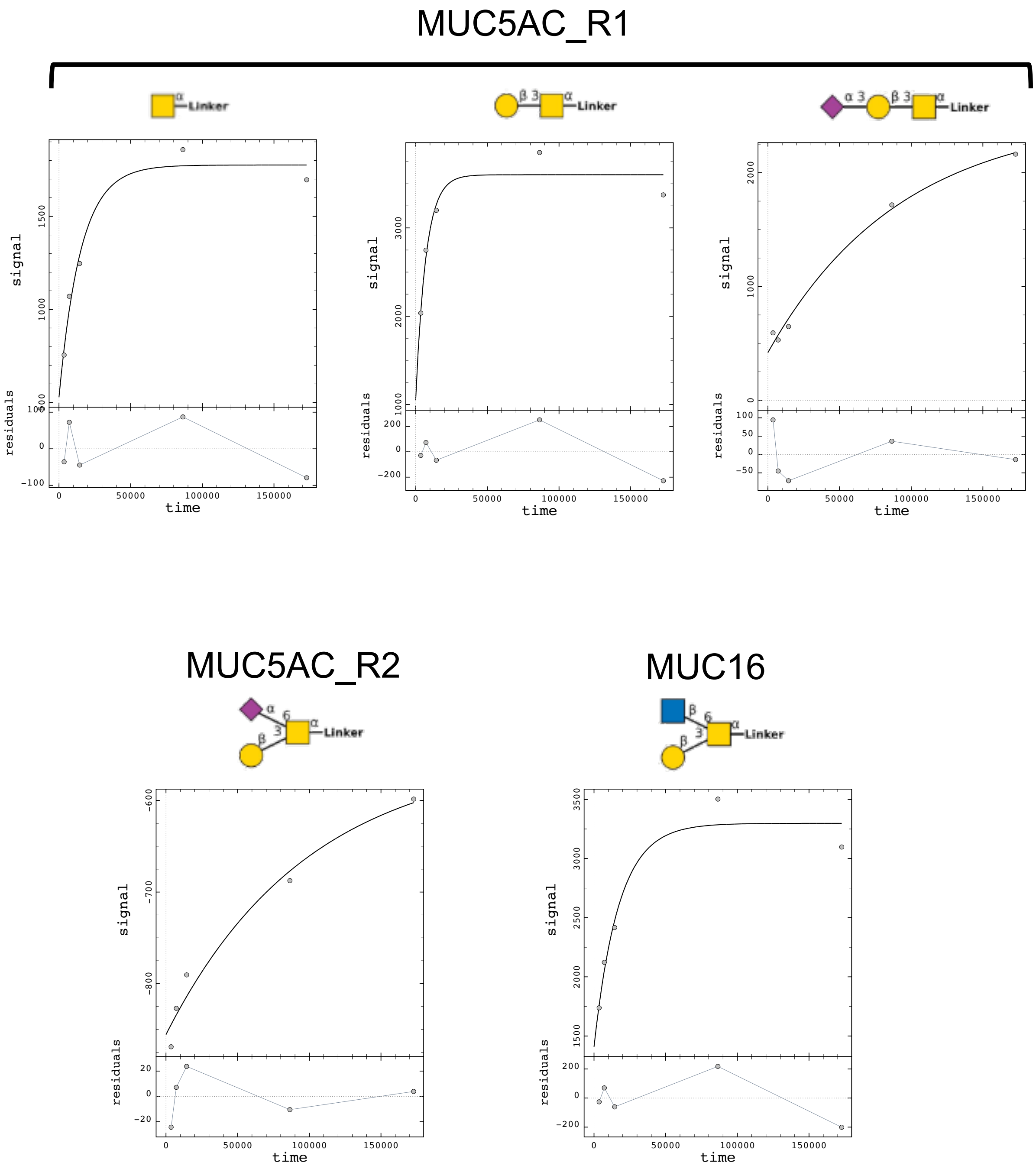

**Supplementary Figure 4.** Examples of FRET screen data converted to fluorescence difference values and fit to a simple bi-molecular kinetic model (*i.e.*, the so-called "hit-and-run" approach). All data shown is for BcM60K.

BcM60K
MUC5AC_R2 Tn

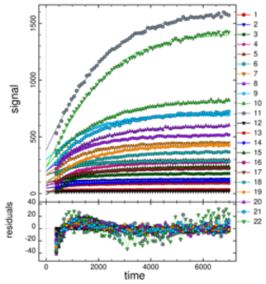

BcM60K
MUC5AC_R2 C1

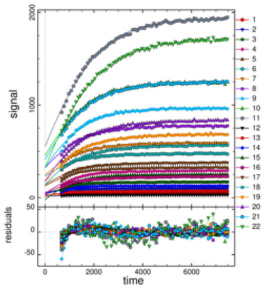

BcM60K
MUC5AC_R2 3SC1

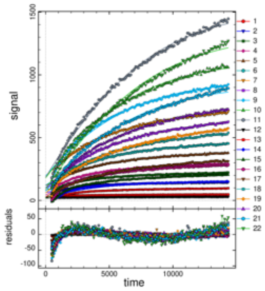

BcM60K
MUC5AC_R2 C2

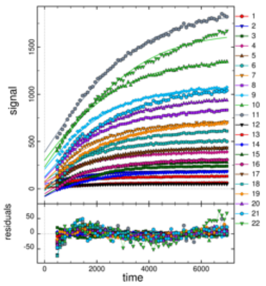

BcM60K
MUC5AC_R2 6SC1

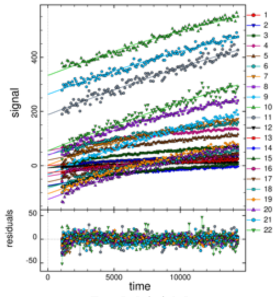

BcM60K
MUC5AC_R2 (repeat)

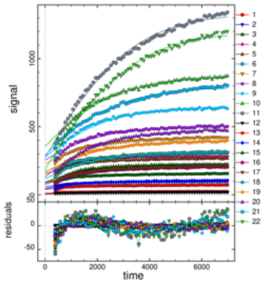

BcM60K
5AC_R2/11 hybrid

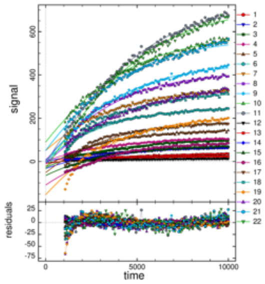

BcM60K
11/5AC_R2 hybrid

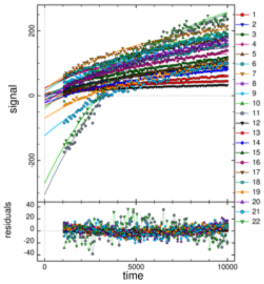

BcM60K
5AC_R2/IgA hybrid

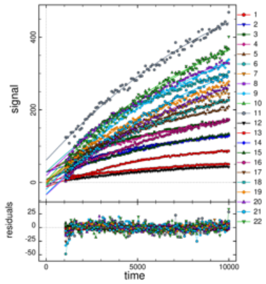

BcM60K
4/5AC_R2 hybrid

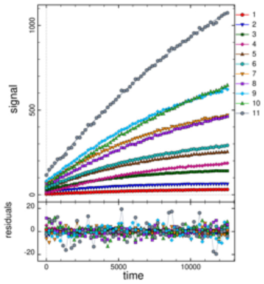

BcM60K
5AC_R2/4 hybrid

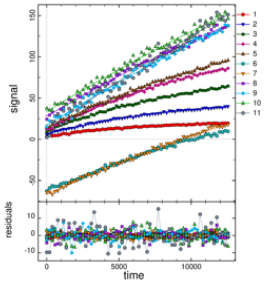

BcM60C
MUC5AC_R2 Tn

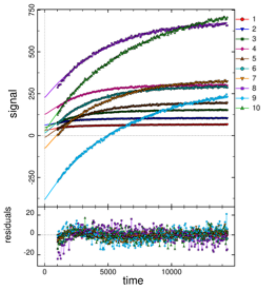

BcM60C
MUC5AC_R2 C1

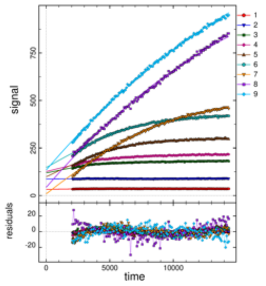

BcM60F
MUC5AC_R2 Tn

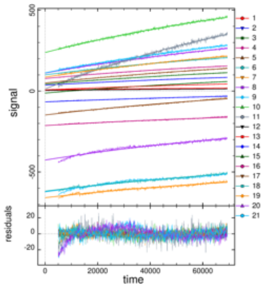

BcM60F
MUC5AC_R2 C1

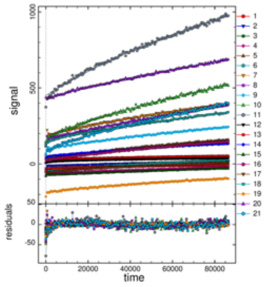

BcM60F
MUC5AC_R2 3SC1

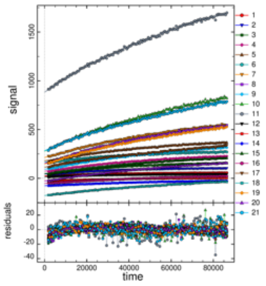

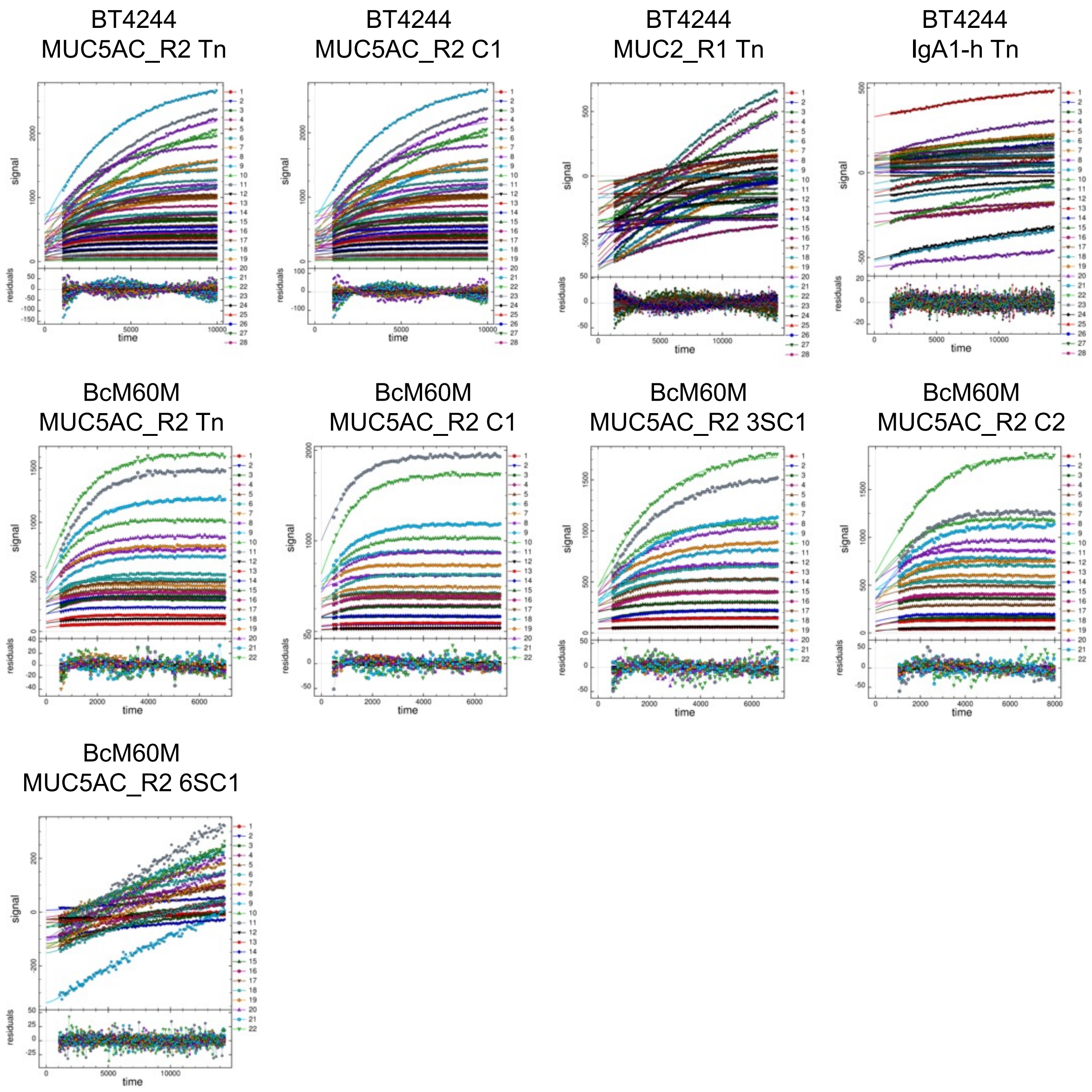

**Supplementary Figure 5 (and preceding page).** Enzyme progress curves used for deriving kinetic constants. The enzyme and substrate used are shown. Symbols indicate the experimental data and solid lines the fit to the model as described in the materials and methods.

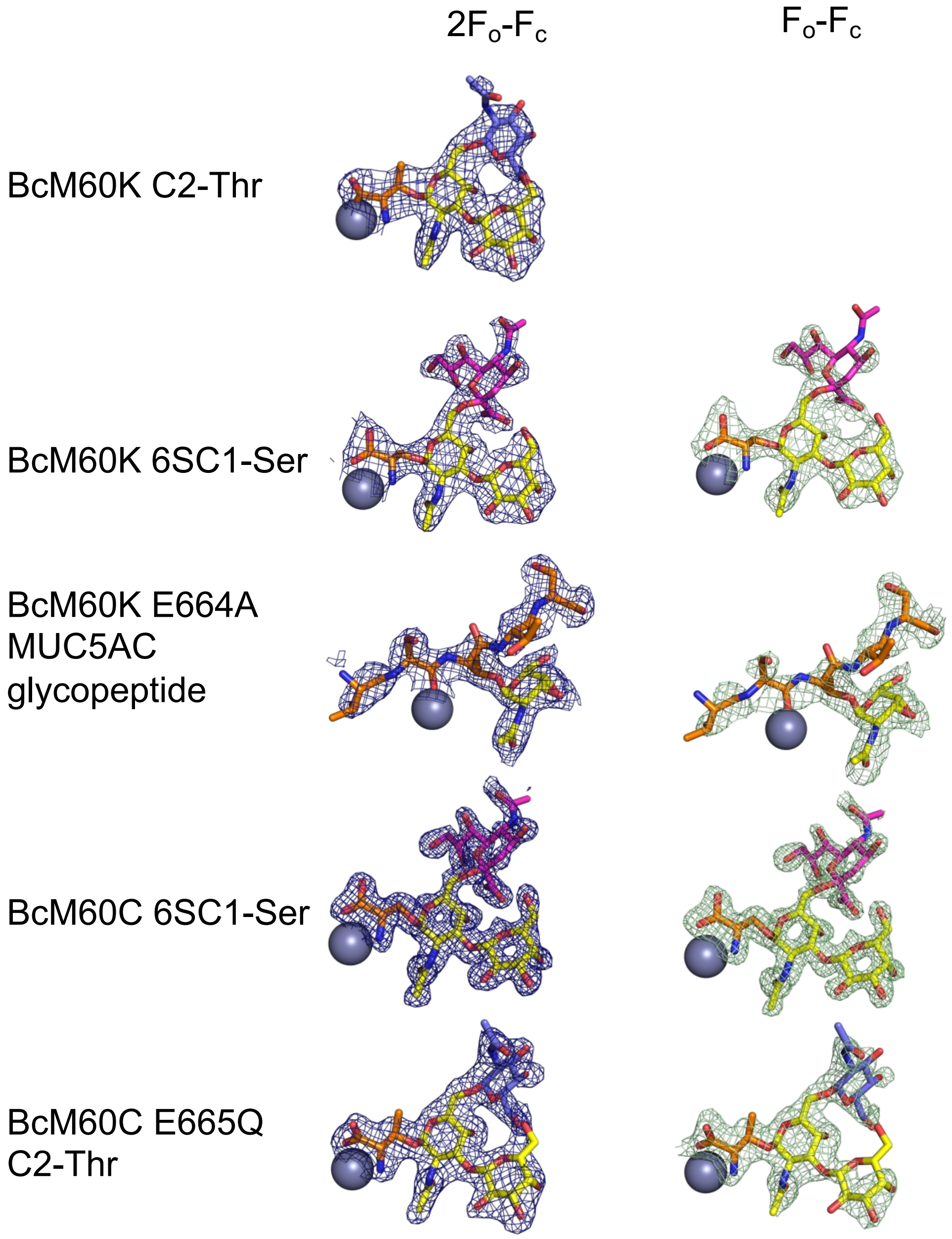

**Supplementary Figure 6.** Experimental electron density maps for ligands in the BcM60K and BcM60C complexes. $2F_o$-$F_c$ electron density maps are contoured at 1$\sigma$ and shown as blue mesh. $F_o$-$F_c$ electron density maps generated by refinement in the absence of ligand are contoured at 3$\sigma$ and shown as green mesh. The $2F_o$-$F_c$ and $F_o$-$F_c$ electron density maps for the BcM60K E664A peptide complex are shown contoured at 0.9$\sigma$ and 2.5$\sigma$, respectively.

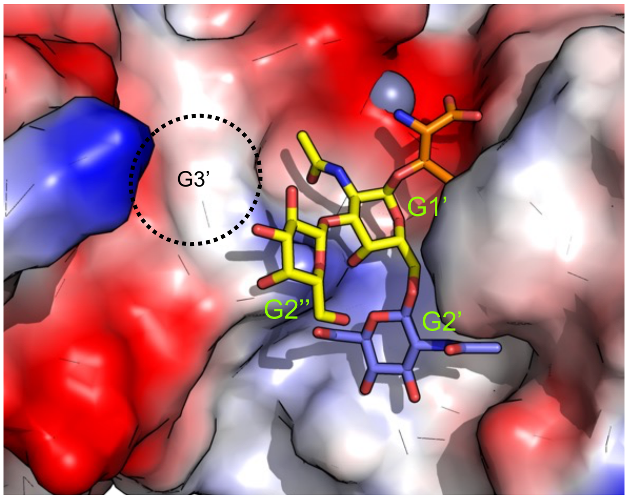

**Supplementary Figure 7.** The surface of the active site of a BcM60K model generated with AlphaFold 3. The C2-Thr was modeled in based on an overlap with the BcM60C C2-Thr complex.

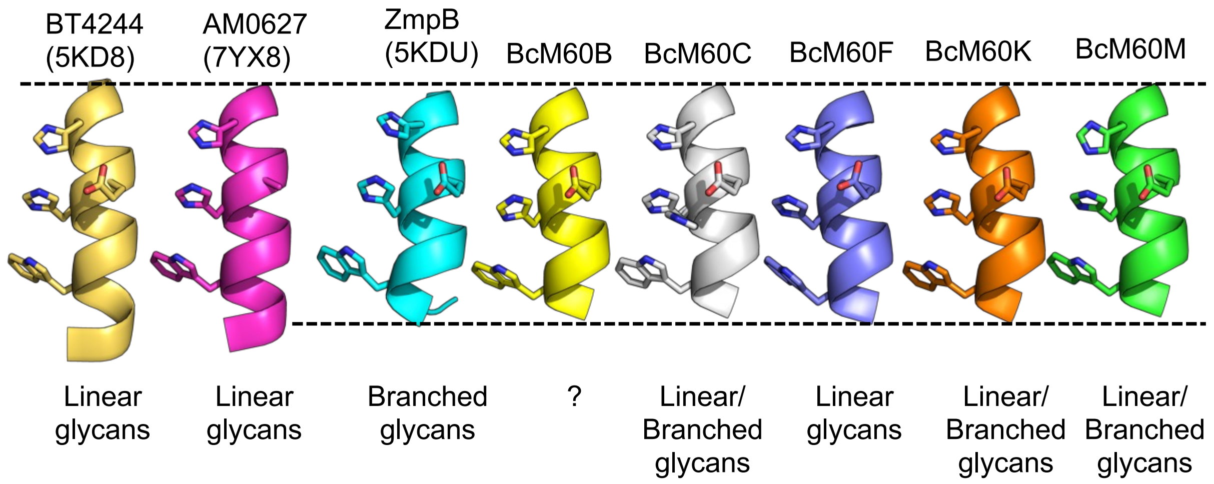

**Supplementary Figure 8.** Comparison of catalytic helix length in characterized peptidase_M60 O-glycopeptidases and in the new BcM60 enzymes. The zinc binding histidines, catalytic glutamate, and GalNAc binding tryptophan are shown for reference. AM0627 refers to AMUC_0627; this structure was an alanine mutant of the catalytic glutamate.

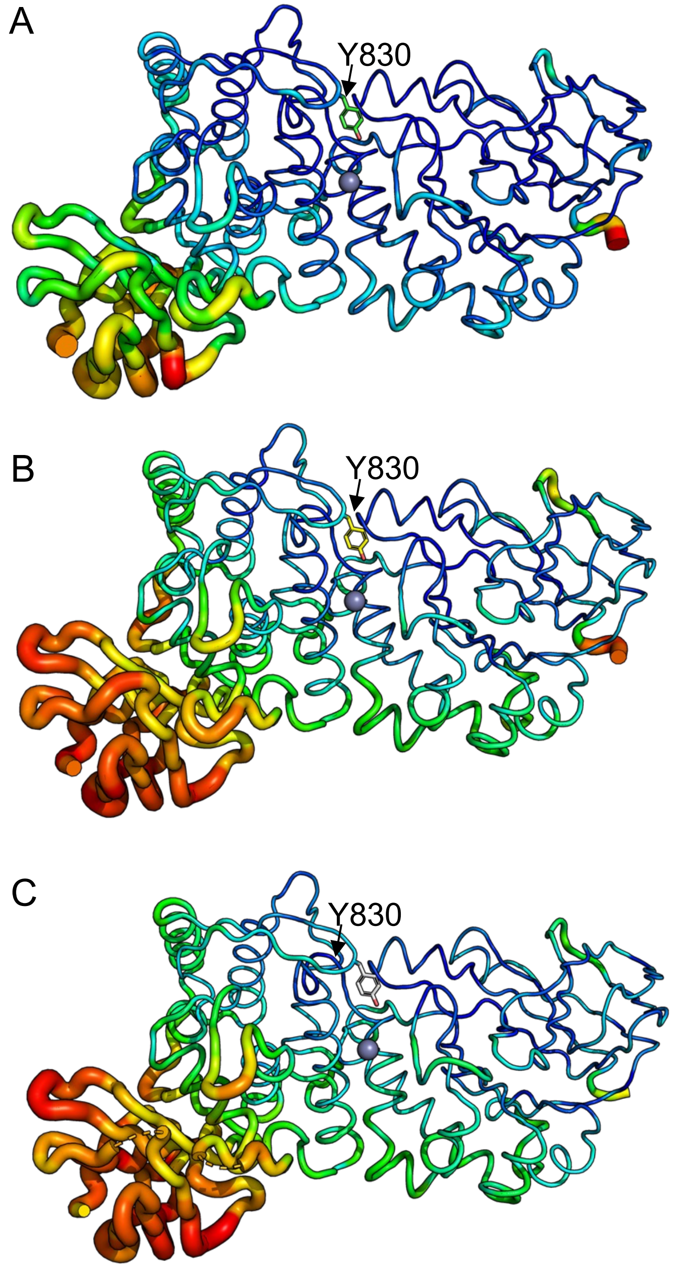

**Supplementary Figure 9.** Tube representations of BcM60K structures. A) complex of BcM60K E664A with the MUC5AC glycopeptide. B) Complex of BcM60K with 6SC1-Ser. C) Unliganded BcM60K. The thickness of the tube from thin to thick and color ramp from blue to red indicate increasing B-factors. The tyrosine residue closing in on the active site is shown as sticks.

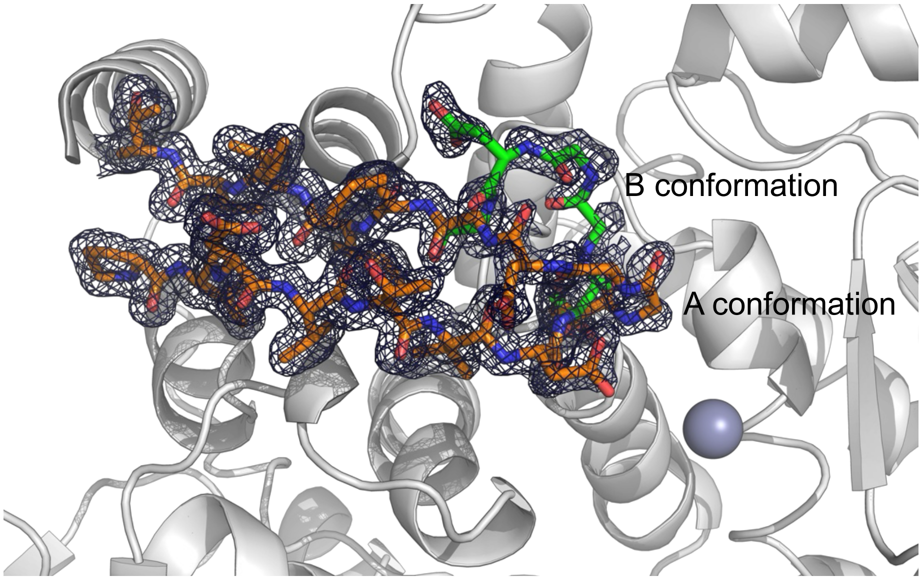

**Supplementary Figure 10.** Dual conformations of the active site loop in BcM60C. The main conformation is shown as orange sticks and the minor, retracted conformation as green sticks. The $2F_o$-$F_c$ electron density map contoured at $1\sigma$ is shown as a mesh.

| | | | P3 | P2 | P1 | P1' | P2' | P3' | P4' | | | IsoGlycP score | Overall Activity Rating |
|---|---|---|---|---|---|---|---|---|---|---|---|---|---|
| MUC5AC_R1 | D | V | K | P | P | T | T | S | T | T | S | 5.5 | Good |
| | P | T | T | S | T | T | S | A | P | P | V | 4.0 | |
| MUC5AC_R2 | P | V | P | A | T | T | V | G | P | V | P | 11.5 | Good |
| MUC5B_R1 | D | V | K | G | S | T | A | T | P | S | S | 7.4 | Good |
| | K | G | S | T | A | T | P | S | S | T | P | 5.0 | |
| | A | T | P | S | S | T | P | G | V | S | K | 10.1 | |
| MUC5B_R2 | T | T | T | A | T | T | P | T | P | V | S | 6.6 | Good |
| | T | A | T | T | P | T | P | V | S | K | G | 9.6 | |
| MUC7_R1 | T | D | V | K | T | T | A | A | P | P | T | 4.0 | Good |
| | T | A | A | P | P | T | P | S | A | T | V | 13.3 | |
| | P | T | P | S | A | T | V | S | K | G | E | 2.7 | |
| MUC16 | V | K | S | V | P | T | T | S | T | P | G | 4.5 | Good |
| | K | S | V | P | T | T | S | T | P | G | T | 4.8 | |
| | V | P | T | T | S | T | P | G | T | S | - | 4.9 | |
| MUC17_R1 | D | V | K | S | P | T | N | S | S | P | T | 3.9 | Good |
| | T | N | S | S | P | T | T | A | E | V | S | 3.8 | |
| MUC1_R1 | P | A | P | G | S | T | A | P | P | A | H | 16.9 | Medium |
| MUC2_R3 | K | S | P | P | P | T | S | T | T | T | L | 8.0 | Medium |
| | P | P | T | S | T | T | T | L | P | V | S | 2.0 | |
| MUC8_R2 | V | H | E | L | P | T | S | S | P | G | G | 7.6 | Medium |
| MUC10 | T | T | D | S | T | T | P | A | P | T | T | 8.4 | Medium |
| MUC11 | T | T | P | A | P | T | T | V | S | K | G | 8.2 | Medium |
| 4/5AC_R2 | S | T | G | H | A | T | V | G | P | V | P | 4.3 | Medium |
| 5AC_R2/4 | P | V | P | A | T | T | P | L | P | V | T | 8.2 | Medium |
| 5AC_R2/11 | P | V | P | A | T | T | H | T | T | L | S | 1.0 | Medium |
| MUC2_R2 | V | K | V | P | P | T | T | T | P | S | P | 20.2 | Poor |
| | V | P | P | T | T | T | P | S | P | P | P | 16.1 | |
| MUC7_R2 | V | K | P | S | A | T | T | P | A | P | P | 3.2 | Poor |
| | K | P | S | A | T | T | P | A | P | P | S | 8.8 | |
| MUC11/12 | S | S | P | G | S | T | H | T | T | L | S | 2.3 | Poor |
| ASF | K | A | E | A | P | T | A | V | P | D | K | 8.4 | Poor |
| Glycoprotein Iba | P | A | P | S | P | T | T | P | E | P | T | 6.6 | Poor |
| | A | P | S | P | T | T | P | E | P | T | V | 6.8 | |
| | T | T | P | E | P | T | V | S | K | G | E | 5.4 | |
| MUC2_R1 | V | K | T | T | V | T | P | T | P | T | P | 5.0 | Poor |
| | T | T | V | T | P | T | P | T | P | T | G | 46.0 | |
| | V | T | P | T | P | T | P | T | G | V | S | 20.5 | |
| | P | T | P | T | P | T | G | V | S | K | G | 5.7 | |
| 11/5AC_R2 | S | S | P | G | S | T | V | G | P | V | P | 26.6 | Poor |
| 5AC_R2/IgA1 | P | V | P | A | T | T | P | S | P | S | P | 16.4 | Poor |
| IgA1/5AC_R2 | P | S | T | P | P | T | V | G | P | V | V | 25.1 | Poor |
| MUC1_R2 | K | A | H | G | V | T | S | A | P | D | T | 3.3 | NA |
| IgA1_h | P | S | T | P | P | T | P | S | P | S | V | 35.9 | NA |
| MUC4 | S | T | G | H | A | T | P | L | P | V | T | 3.0 | NA |
| MUC3A/B | S | S | I | T | T | T | E | T | T | S | V | 1.8 | NA |
| MUC8_R1 | P | L | Q | E | G | T | P | G | S | R | V | 3.0 | NA |
| MUC19 | D | V | K | S | T | T | V | A | P | G | S | 5.2 | NA |
| MUC20 | V | A | P | G | S | T | T | V | V | S | K | 4.3 | NA |
| MUC6_R1 | T | T | T | Y | P | T | P | S | H | P | Q | 6.9 | NA |

**Supplementary Figure 11.** Summarized activity of BcM60K and BcM60C based on the FRET screen analysis. Likelihood of glycosylation was estimated by IsoGlyP with higher scores indicating higher likelihood of begin an acceptor site for *O*-glycosylation by ppGalNAcT2.
